# The neuropeptide FLP-17 regulates an oviposition behavior in the nematode *Caenorhabditis elegans* that increases maternal reproductive fitness in low oxygen environments

**DOI:** 10.1101/2022.11.15.516679

**Authors:** Tong Young Lee, Eunha Chang, Kyoung-hye Yoon, Jin I. Lee

## Abstract

**SUMMARY:** The ability for animals to adapt their behaviors to specific environments is imperative to increase their evolutionary success. This is particularly true for behaviors such as parental behaviors that directly affect reproductive fitness. Here, we identified an oviposition behavior in the nematode *C. elegans* that increases the survival of the young. In standard laboratory culture, the bacterivorous hermaphrodite mothers lay eggs across a 2D *E. coli* lawn with no discernable pattern. However, in 3D culture they display a stereotypical behavior in which they temporarily leave the bacteria to lay eggs far away from the *E. coli* colony, resulting in a scattered ring of eggs located outside the bacteria. This oviposition behavior requires low oxygen levels and is regulated by the neuropeptide FLP-17 and its cognate receptor EGL-6. We confirm that a circuitry involving the oxygen-sensing BAG neurons and the vulva muscle-controlling HSN motor neurons regulates oviposition behavior. We show that loss of proper oviposition behavior results in lower reproductive fitness for the mothers and embryonic lethality for the eggs laid in bacteria under hypoxic conditions. Finally, we show that the degree of oviposition behavior varies among wild strains of *C. elegans* found in nature. The ability for *C. elegans* mothers to sense their environments and adjust their behaviors in adverse conditions is likely an adaptation that has allowed the worm to thrive in diverse and often hazardous habitats.

## INTRODUCTION

Parental behaviors have a direct influence on the survival of offspring, contributing to the reproductive fitness of the parent. The success of the behaviors, however, are subject to the environment and the ecological niche that the animal inhabits. Mothers must adapt their maternal behaviors to unstable and often dangerous conditions^1^. This type of behavioral plasticity in parental behaviors is seen in response to ecological factors such as food availability in sparrows^2^, to predators in various birds^3^, and to increasing temperature in the European great tit^4^. In particular, rising temperatures due to climate change have resulted in a directed selection of earlier egg laying in great tit mothers, demonstrating that a genetic component is likely evolving due to such environmental changes^4^.

Studies in genetic model animals such as mice and fruit flies have revealed specific genes and neurons that regulate parental behaviors^5-8^. Moreover, *Drosophila melanogaster* displays an oviposition behavior in which mother flies, guided by sensory cues, lay their eggs in specific locations in both the laboratory^8-10^ and in nature^11^. In fact, mothers can vary their oviposition behavior depending on environmental context^8^. Whether such behavioral plasticity of oviposition behavior confers reproductive fitness benefits to the mothers is not known.

The behaviors of the free-living bacterivorous nematode *C. elegans* have been studied in the laboratory extensively. The worm can sense and respond to many environmental cues such as odors and pheromones, temperature, humidity and gases such as oxygen and carbon dioxide^12^. In normal laboratory culture, *C. elegans* hermaphrodite mothers do not appear to display any specific oviposition behaviors, laying eggs across the bacterial lawn they feed on without any discernable pattern^13^. However, *C. elegans* is endowed with a relatively sophisticated egg-laying circuitry consisting of over a dozen uterine and vulval muscles, several motor neurons, and two neurotransmitters, which allow exquisite control of its vulva muscles during egg laying behavior.

*C. elegans* in nature can be found thriving embedded in fruit and vegetative compost^14^, a much more complex three-dimensional environment than the 2D controlled conditions in the laboratory. Such microbe and predator rich habitats can pose challenges and dangers for the worm to survive and navigate, particularly for smaller and more vulnerable *C. elegans* larvae^15,16^.

Previously, we designed 3D habitats called Nematode Growth Tube (NGT-3D) and Nematode Growth Bottle (NGB-3D) in which *C. elegans* and their laboratory food *E. coli* bacteria are embedded in an agar media^17,18^. Compared with the normal laboratory 2D culture plates, these habitats better reflect the complex 3D spatial environment that *C. elegans* is exposed to in nature.

In this study, we identify a novel oviposition behavior that *C. elegans* mothers only display in the 3D environment. We find that wild-type mothers shift the location of egg laying specifically in a low oxygen environment that is common when bacterial colonies are embedded in media. The plasticity of oviposition behavior is mediated by the neuropeptide FLP-17 and its cognate receptor EGL-6 that inhibits egg laying. We demonstrate that *flp-17* mutant mothers defective in oviposition behavior display decreased reproductive fitness in 3D. Finally, we test genetically diverse wild *C. elegans* strains and found a large degree of variability in oviposition behavior.

This demonstrates genetically that the plasticity of maternal behaviors could be an adaptive product of evolution.

## RESULTS

### A novel oviposition behavior by *C. elegans* in a three-dimensional environment

Previously, we designed NGB-3D (Fig 1B) and NGT-3D (Fig 3A) habitats that better reflect the 3D soil and rotting vegetation and fruit environments that *C. elegans* is often found in^14,17,18^. In 2D culture, *E. coli* strain OP50 bacteria is spread out and grows in a flat lawn that *C. elegans* inhabits (Fig 1A), whereas in 3D the bacteria grow as small dense colonies embedded in the agar (Fig 1B), and worms develop, reproduce normally and have normal lifespans in these 3D environments^17^. Despite their normal growth in 3D, we previously demonstrated that the full sensory capabilities of *C. elegans* were required for the worms to survive in a 3D environment. A mutant strain with compromised sensory capabilities was able to survive normally in the 2D NGM environment but could not survive in the 3D environment^17^. Thus, we surmised that sensory behaviors that were hidden in flat 2D plates may be revealed in our 3D environments.

**Figure 1.**
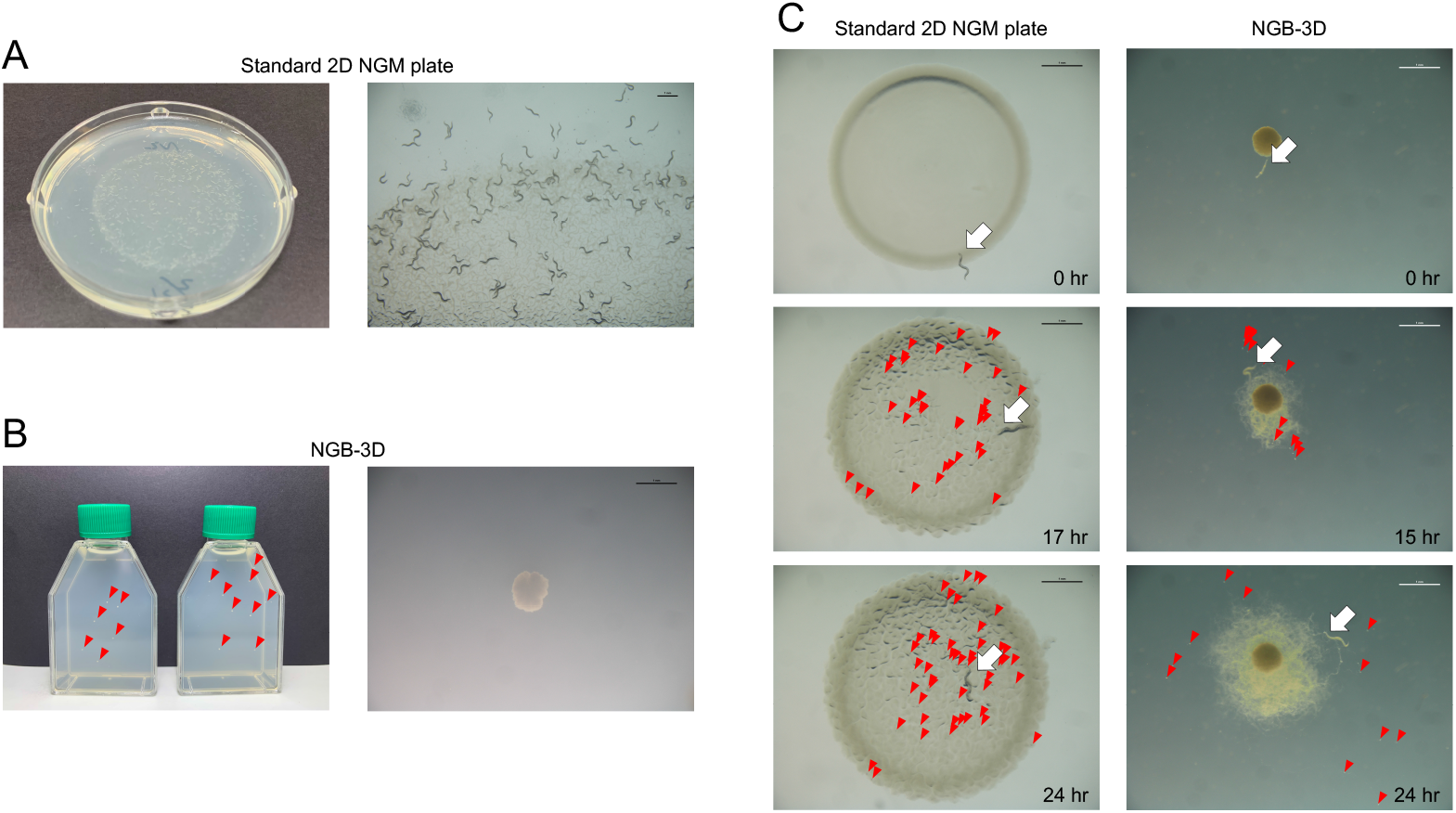
*C. elegans* behavior in 2D and 3D environments. **(A)** Standard laboratory 2D NGM cultivation plate (left). Close-up showing *C. elegans* adult worms (right) Scale bar indicates 1mm.. (B) NGB-3D nematode cultivation bottles. Red arrowheads indicate *E. coli* OP50 strain colonies. Close-up of an *E. coli* colony. Scale bar indicates 1mm. (C) Behavioral differences between 2D and 3D environment. An 2D NGM plate (left) and NGB-3D (right) were imaged for 24 hrs. White arrow indicates location of *C. elegans* mother. Red arrows indicate eggs laid. Scale bar indicates 1mm.

To compare the behaviors in the two environments, we placed single hermaphrodite mothers either on a flat 2D NGM plate or NGB-3D and imaged the worms for 24 hrs (Fig 1C; Movie S1, Movie S2). We observed three major differences in behavior: 1) In 2D the mother worm preferred to stay inside the bacteria lawn, whereas in 3D the mother mostly stayed outside the bacteria colony, while keeping its head in the bacteria (Fig 1C, white arrow); 2) In 2D the mother worms moved randomly within the bacteria lawn, whereas in 3D the mother spread the bacteria outwardly from the colony in a “sprawling” fashion (Fig 1C); 3) in 2D the mother worm laid eggs inside the bacteria lawn haphazardly, whereas in 3D the mother laid eggs away from the bacteria, surrounding the colony with her eggs (Fig 1C, red arrows).

To quantify these observations, we measured the average location of the mother worms in each colony in relation to the center of the bacterial lawn or sprawl. We found that, in 2D NGM, nearly 80% of adult hermaphrodite mothers remain inside the bacterial lawn, whereas in 3D, worms prefer to stay towards the edge of the bacterial sprawl, occasionally venturing inside the colony (Fig 2A, B). *C. elegans* navigates its environment by displaying short forward and short reversal movement, as well as long straight runs and “omega” turns in which worms change their direction around at least 90 degrees^19^. Comparing the navigation movements of the mother in 2D and 3D, we found that the movements of the mother worms did not alter dramatically in 3D (Fig S1), indicating that in 2D and 3D navigation behaviors are similar.

**Figure 2.**
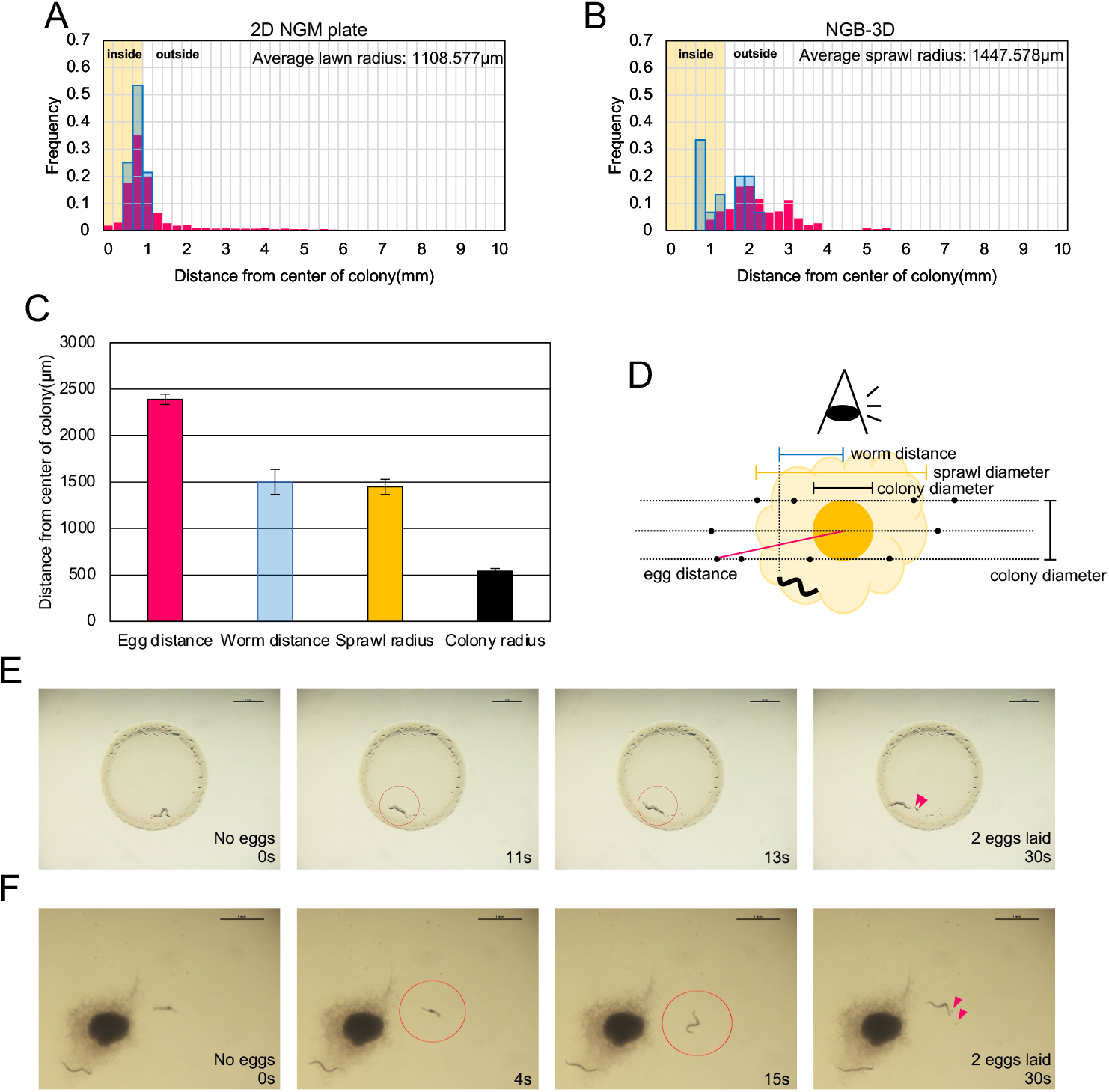
*C. elegans* mothers display an oviposition behavior in a 3D environment. (A-B) Spatial distribution of laid eggs (red) and location of mother (blue) in 2D NGM plate (A) and NGB-3D (B). Yellow shading indicates area inside the bacteria. (C-D) Colony and sprawl size, worm distance and egg distance in 3D (C) and measurement method in 3D (D). Average worm and egg distance was calculated by averaging all mother *C. elegans* and eggs inhabiting a single E. coli colony. Error bars indicate standard error. (E-F) 30-second video time lapse images of egg laying of N2 mother in 2D NGM plate (E) and NGB-3D (F). Red circles indicate position moment of egg laying. Red arrowheads indicate laid eggs. Scale bar indicates 1 mm.

We then quantified the differences in egg laying behavior in 2D and 3D by measuring the locations of the eggs in relation to the center of the bacterial lawn or sprawl (Fig 2C, 2D). In the 2D NGM plate most of the eggs were laid inside the bacterial lawn and dispersed throughout the lawn more or less randomly, similar to the locations of the mother worms (Fig 2A, 2C). However, in 3D the majority of eggs were laid far away from the sprawl rather than inside the bacteria (Fig 2B), an average of 2.39 mm away from the center of the colony, and over 800 µm from the edge of the sprawl, far away from where the mother worm prefers to stay (Fig 2C, 2D). We found these distances to be relatively consistent from worm to worm and colony to colony. Thus, unlike the mothers in 2D plates that passively lay eggs on the bacterial lawn (Fig 2E; Movie S3), the hermaphrodite mother in 3D must be actively moving away from the edge of the bacterial sprawl to lay her eggs and then coming back to the bacteria, which we have observed (Fig 2F; Movie S4). This novel behavior can be described using the 3D oviposition ratio, calculated as mother distance from the center of the colony/egg distance from the center of the colony which represents the distance that mothers travel to lay their eggs in 3D (Fig 3I). Oviposition ratio for wild type mother worms is 79%. Thus, we discovered a potential oviposition behavior in 3D environments, which may be a maternal adaptation to maximize the survival and fitness of the young in *C. elegans* native environments.

**Figure 3.**
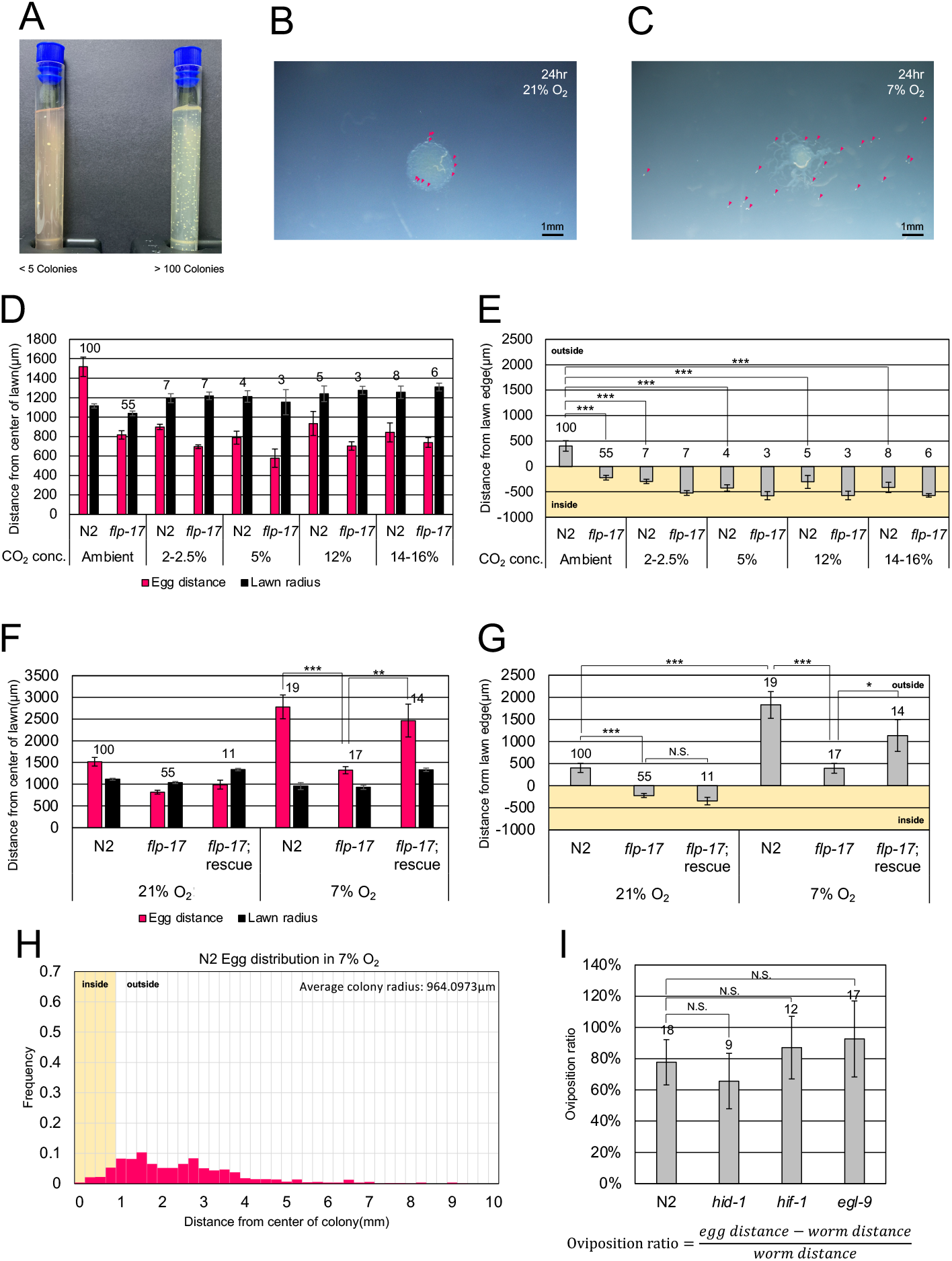
Low oxygen induces oviposition behavior. (A) Increased bacterial colonies decreases oxygen levels in NGT-3D indicated by thioglycolate media color change from pink to clear. (B) Eggs laid after 24 hrs in 2D NGM plate at 21% oxygen. Red arrowheads indicate eggs. Scale bar indicates 1 mm. (C) Eggs laid after 24 hrs in 2D NGM plate at 7% oxygen. Red arrowheads indicate eggs. Scale bar indicates 1 mm. (D) High CO_2_ levels induces *flp-17*-independent egg laying inside the bacteria lawn. Egg distance was calculated as an average of all eggs around each separate lawn. The number of lawns counted is indicated above the bars. Error bars indicate standard error. (E) Distance between lawn edge and egg location in high CO_2_. Statistical significance by student’s T-test is indicated. Error bars indicate standard error. (F) Low O_2_ levels induces a *flp-*17-dependent oviposition behavior away from the bacteria. Egg distance was calculated as an average of all eggs around each separate lawn. The number of lawns counted is indicated above the bars. Statistical significance by student’s T-test is indicated. Error bars indicate standard error. (G) Distance between lawn edge and egg location in low O_2_. Statistical significance by student’s T-test indicated. Error bars indicate standard error. (H) Spatial distribution of laid eggs in 7% oxygen in 2D in N2 mothers. (I) 3D oviposition ratio in N2 and mutants defective in high CO_2_ and low O_2_ response. Statistical significance by student’s T-test indicated. Error bars indicate standard error. * indicates p < 0.05; ** indicates p < 0.01; *** indicates p < 0.001; NS indicates not significant.

### Low oxygen induces an oviposition behavior

Since *C. elegans* mothers do not display an oviposition behavior in the 2D environment, we surmised that some change unique to the NGB-3D environment must be sensed by mothers to induce egg laying far away from the bacteria. Since the mother worm and *E. coli* colonies are embedded in agar, baseline oxygen concentrations should be low in our 3D environments, and the presence of dense colonies of respiring bacteria could further exacerbate gas conditions, increasing CO_2_ levels while decreasing O_2_ levels. Using our NGT-3D test tubes, we grew OP50 *E. coli* colonies in thioglycolate medium that changes from a pink color in oxygenated conditions to a clear color in anoxic conditions^20,21^. Oxygen levels were clearly lower when more bacterial colonies were present in 3D (Fig 3A).

Facultative anaerobic *E. coli* bacterial colonies can alter both carbon dioxide and oxygen surrounding levels. To further delineate whether CO_2_ or O_2_ conditions contributes to oviposition behavior, we wanted to alter gas levels and observe any changes in egg laying. Since we were unable to control internal gas concentrations in our 3D environments where *C. elegans* and bacteria are embedded in the media, we instead used a setup in which elements of 3D conditions can be replicated in a 2D environment (Fig S2). Previously, we showed that some elements of 3D behavior can be replicated by using 2D “dot” plates with tiny (∼1mm radius) bacterial lawns^17^. These plates could be placed in a closed chamber to control 2 CO_2_ or O_2_ levels, allowing us to see the effect of different gas concentrations on oviposition behavior.

As observed before, we found that in ambient air conditions, wild-type mothers lay eggs both inside and occasionally just outside of the colony, but in general stay close to the bacteria (Fig 3B, 3D, 2A). When we increased ambient CO_2_ concentration, in all concentrations tested, mothers rarely laid eggs outside of the lawn (Fig 3D-3E). Thus, high CO_2_ does induce a mild change in behavior, but opposite of what we observe in 3D.

Next, we wondered if changes in oxygen level could induce an oviposition behavior. Remarkably, at 7% O_2_ we found that wild-type mothers tended to lay their eggs far away from the bacteria, similar to what we observe in NGB-3D (Fig 3C, 3F-3H, Fig S3). Whereas in normal oxygen mother worms lay nearly 80% of their eggs inside the bacteria (Fig 2C), in 7% oxygen more than 90% of the eggs were laid outside, with nearly 50% of the eggs laid more than 2 mm from the center of the bacterial lawn (Fig 3H), resembling what we see in NGB-3D egg laying (Fig 2D). Thus, low oxygen drives the mothers to display oviposition behavior.

To investigate how gas levels induce *C. elegans* 3D egg laying behavior, we assessed 3D oviposition behavior in mutants of genes that function in high CO_2_ or low O_2_ environments. HID-1, a component of neuropeptide signaling, is necessary for proper response to acute carbon dioxide^22^. However, *hid-1* mutant mothers that are defective for CO_2_ response displayed normal oviposition behavior (Fig 3I). The hypoxia-induced factor HIF-1 along with the prolyl hydroxylase EGL-9 allow *C. elegans* to adapt behaviors to cultivation in hypoxic conditions^23-25^. However, both *hif-1* and *egl-9* mutant mother worms displayed normal oviposition behavior in NGB-3D (Fig 3I).

### FMRF-like neuropeptide FLP-17 and its cognate receptor EGL-6 regulate oviposition behavior

The FLP-17 FMRF-like neuropeptide is known to mediate a low oxygen signal from the BAG sensory neuron to intestinal cells via the EGL-6 receptor^26,27^. In addition, FLP-17 through the EGL-6 receptor is known to inhibit egg laying^28^. We wondered whether FLP-17 could also be regulating 3D oviposition behavior. In the standard 2D NGM plates, *flp-17* mutants remained in the bacterial lawn and the distribution of the laid eggs were similar to N2 wild-type, although on average *flp-17* mothers laid eggs closer to the lawn than N2 (Fig 4A, 4C, and 3D). In NGB-3D, *flp-17* mutant mothers also tended to remain near the edge of the bacterial sprawl (Fig 4B). However, unlike N2, they laid the majority of eggs close to the bacterial sprawl (Fig 2D, Fig 4D). We captured this defective oviposition behavior in video showing *flp-17* mothers laying eggs close to the bacteria where she prefers to reside (Movie S5, Fig 4A). Overall, we found that *flp-17* mutant worms lay eggs 1.54 mm from the center of the colony and only 0.3 mm from the distance where the mother worm is normally located (Fig S3), displaying an oviposition ratio of 32%, in contrast to the oviposition ratio of 79% in wild-type mothers (Fig 4D). Finally, transgenic expression of wild-type *flp-17* behavior, confirming that *flp-17* is sufficient to induce this behavior (Fig 4D and 4F).

**Figure 4.**
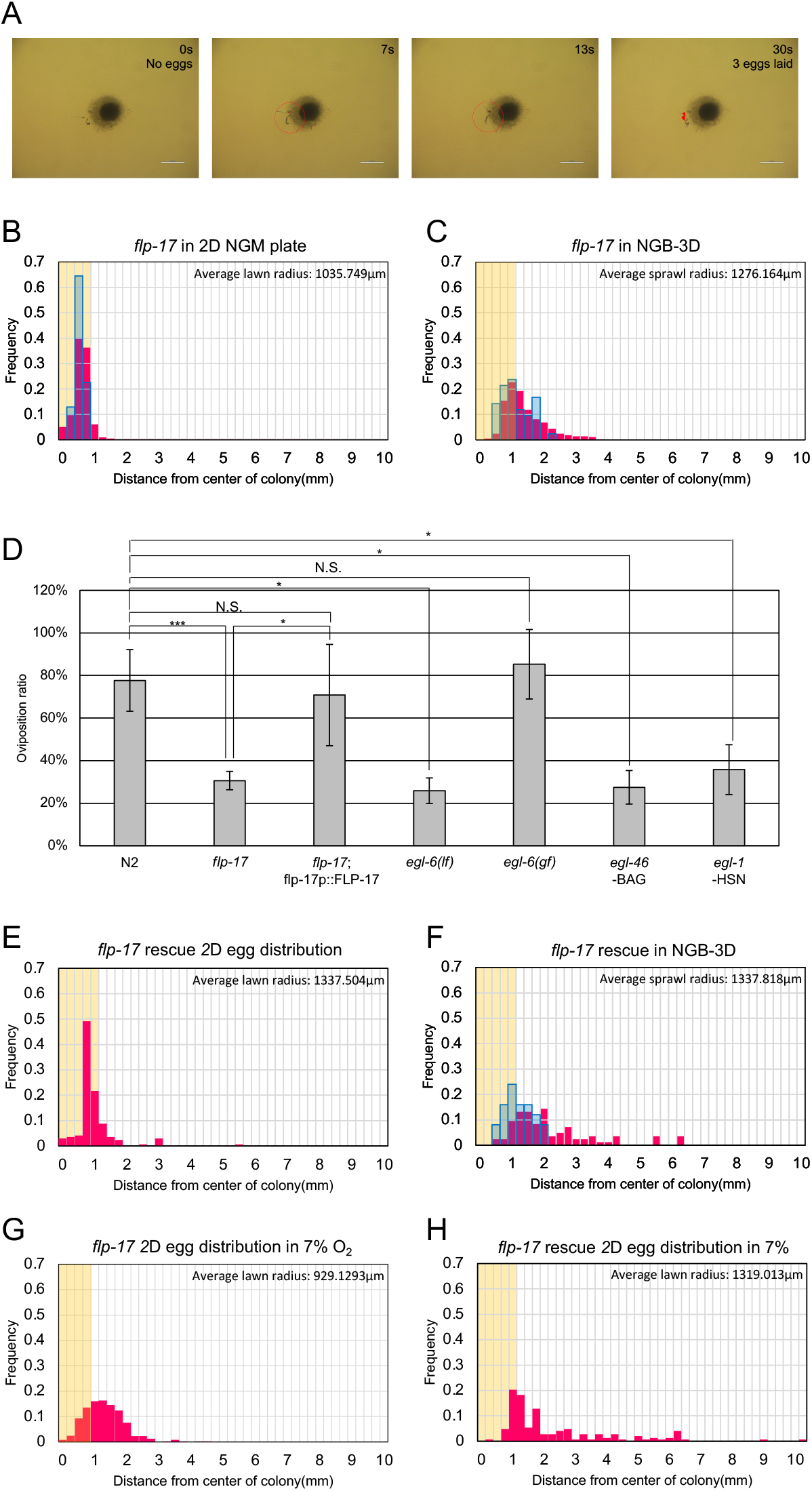
FLP-17 and EGL-6 mediate oviposition behavior through a BAG sensory neuron and HSN motor neuron circuitry. (A) 30-second video time lapse images of egg laying of N2 mother in NGB-3D. Red circle indicates egg laying moment. Red arrow indicates laid eggs. Scale bar indicates 1 mm. (B) *flp-17* spatial distribution of laid eggs and location of mother in 2D NGM plate (C) *flp-17* spatial distribution of laid eggs and location of mother in NGB-3D. Egg distance in 2D NGM in flp-17 mutants in different oxygen conditions. (D) Spatial distribution of laid eggs and location of mother in normal oxygen in 2D in *flp-17* mutants. Statistical significance by student’s T-test is indicated. Error bars indicate standard error. (E) Spatial distribution of laid eggs and location of mother in *flp-17* strains. Spatial distribution of laid eggs and location of mother in NGB-3D in flp-17 rescue strain (F) Spatial distribution of laid eggs in normal and 7% oxygen in 2D in *flp-17* rescue strain. (G) Spatial distribution of laid eggs in 7% oxygen in 2D in *flp-17* mutants. (H) Spatial distribution of laid eggs in 7% oxygen in 2D in *flp-17* rescue strain. * indicates p < 0.05; ** indicates p < 0.01; *** indicates p < 0.001; NS indicates not significant.

FLP-17 neuropeptide is released by the BAG sensory neurons and binds to the G-protein coupled receptor EGL-6 on the HSN motor neurons to inhibit egg laying behavior^28^(Fig 6). We tested whether mutants of *egl-6* were also defective in 3D oviposition behavior. In contrast to a gain-of-function *egl-6* mutant that displayed normal 3D egg laying, loss-of-function *egl-6* mutant mothers were defective in oviposition behavior, similar to *flp-17* mutants (Fig 4D; Fig S3). Thus, we establish that FLP-17 ligand and its cognate receptor EGL-6 can regulate 3D oviposition behavior.

### BAG sensory neurons and the HSN motor neuron are required for proper oviposition behavior

The *flp-17* gene is expressed solely in the BAG sensory neurons that are known to sense O_2_ and CO_2_ gas^29-32^. To establish the role of the BAG sensory neurons in oviposition behavior, we tested 3D egg laying in a mutant strain of *egl-46*, which lacks the BAG neuron due to the loss of a transcription factor required for BAG neuron cell fate^33,34^. We found that *egl-46* mothers also lay their eggs close to the bacteria similar to *flp-17* mutants (Fig 4F; Fig S3).

*C. elegans* egg-laying behavior is regulated by neuromuscular circuitry^13^. The HSN motor neurons control egg laying by activating vulva muscle and is also known to receive input from the BAG sensory neurons^28^. *egl-1* mutants, which lack HSN neurons^35,36^, lay their eggs close to the bacteria similar to *egl-6* mutants (Fig 4F; Fig S3). Taken together, we hypothesize that a BAG to HSN circuitry mediated by FLP-17 and EGL-6 regulate oviposition behavior in our 3D environment.

### FLP-17 regulates low oxygen oviposition behavior

Since FLP-17 from the BAG sensory neurons is necessary for the behavior in 3D, we asked whether FLP-17 also is required for oxygen-dependent oviposition behavior observed in 2D NGM plates. Under normal oxygen conditions, wild-type mothers lay eggs either inside the bacteria or close to the edge of the lawn (Fig 2A, 3B). *flp-17* mutant mothers lay nearly all of their eggs inside the bacteria (Fig 4B). This small difference in *flp-17* egg laying is quantifiable (Fig 3F-3G, Fig S3); however, it is not due to the *flp-17* gene as transgenic rescue of the neuropeptide gene does not restore normal egg laying behavior. Strikingly, at 7% low oxygen conditions, over 90% of the eggs were located either inside the bacteria or within 1000 µm from the edge of bacteria lawn (Fig 3F-3G, 4G, S3), contrary to wild-type mothers whose eggs were widely dispersed (Fig 3H).

As with 3D oviposition behavior, we found that transgenic rescue of *flp-17* restored the low oxygen oviposition behavior (Fig 3F-3G, Fig S3). In addition, we also checked whether *flp-17* mediates the low CO_2_ egg laying behavior we described above (Fig 3D-3E) and found that *flp-17* mutants displayed normal CO_2_-dependent oviposition behavior. Taken together, we demonstrated that FLP-17 mediates a hypoxic oviposition behavior.

### Wild *C. elegans* strains display diverse oxygen-dependent oviposition behavior

An oxygen-dependent oviposition behavior such as the one described should be adapted to the specific environment and ecology that the nematode inhabits. Indeed, varying oxygen sensory behaviors were observed in different wild strains of *C*. elegans^37^. Along with the laboratory N2 strain, we tested 11 genetically diverse wild strains for oxygen-dependent oviposition behavior^38^. In 21% oxygen all the strains displayed similar egg laying behavior, with nearly all eggs laid inside the bacterial lawn (Fig 5A). However, at 7% oxygen even though all the mother worms tended to stay inside the lawn, the 12 strains differed in their oviposition behavior. For instance, strain DL238 mothers laid eggs almost 2 mm from the edge of the lawn similar to N2 mother worms (Fig 5B). In contrast, strains JT11398 and JU258 laid eggs 707 µm and 372 µm from the lawn, respectively, significantly closer to the bacteria than N2 (Fig 5B). This supports the notion that genetic and/or ecological diversity can influence *C. elegans* oviposition behavior.

**Figure 5.**
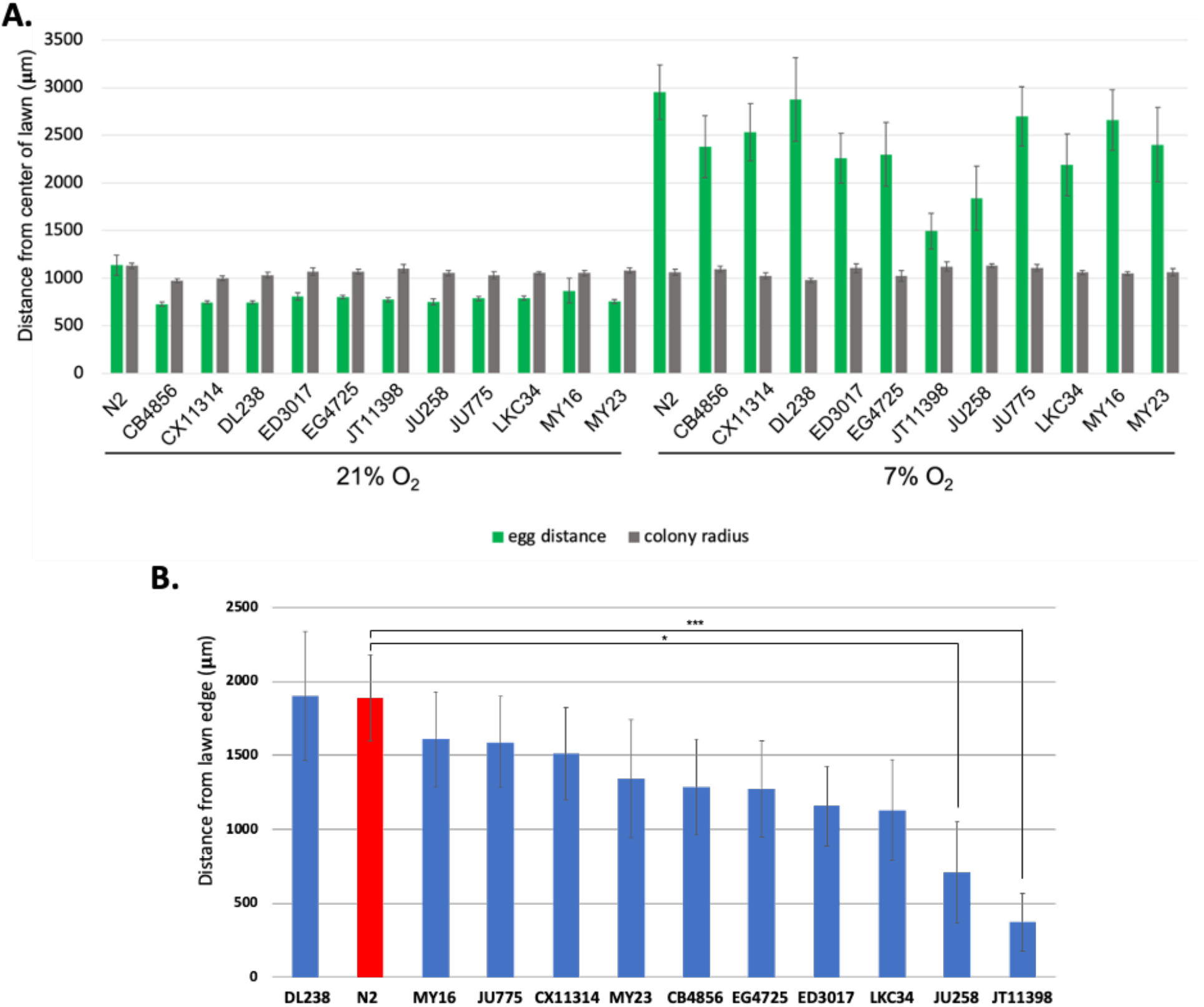
Genetically diverse wild strains of *C. elegans* display variable oxygen-dependent oviposition behavior. (A) Different natural strains of *C. elegans* display different oviposition behavior in low O_2_ levels. Egg distance was calculated as an average of all eggs around each separate lawn. Error bars indicate standard error. Egg distance was calculated as an average of all eggs around each separate lawn. (B) Distance between lawn edge and egg location in 7% O_2_. Statistical significance by student’s T-test is indicated. Error bars indicate standard error. * indicates p < 0.05; ** indicates p < 0.01; *** indicates p < 0.001; NS indicates not significant.

**Figure 6.**
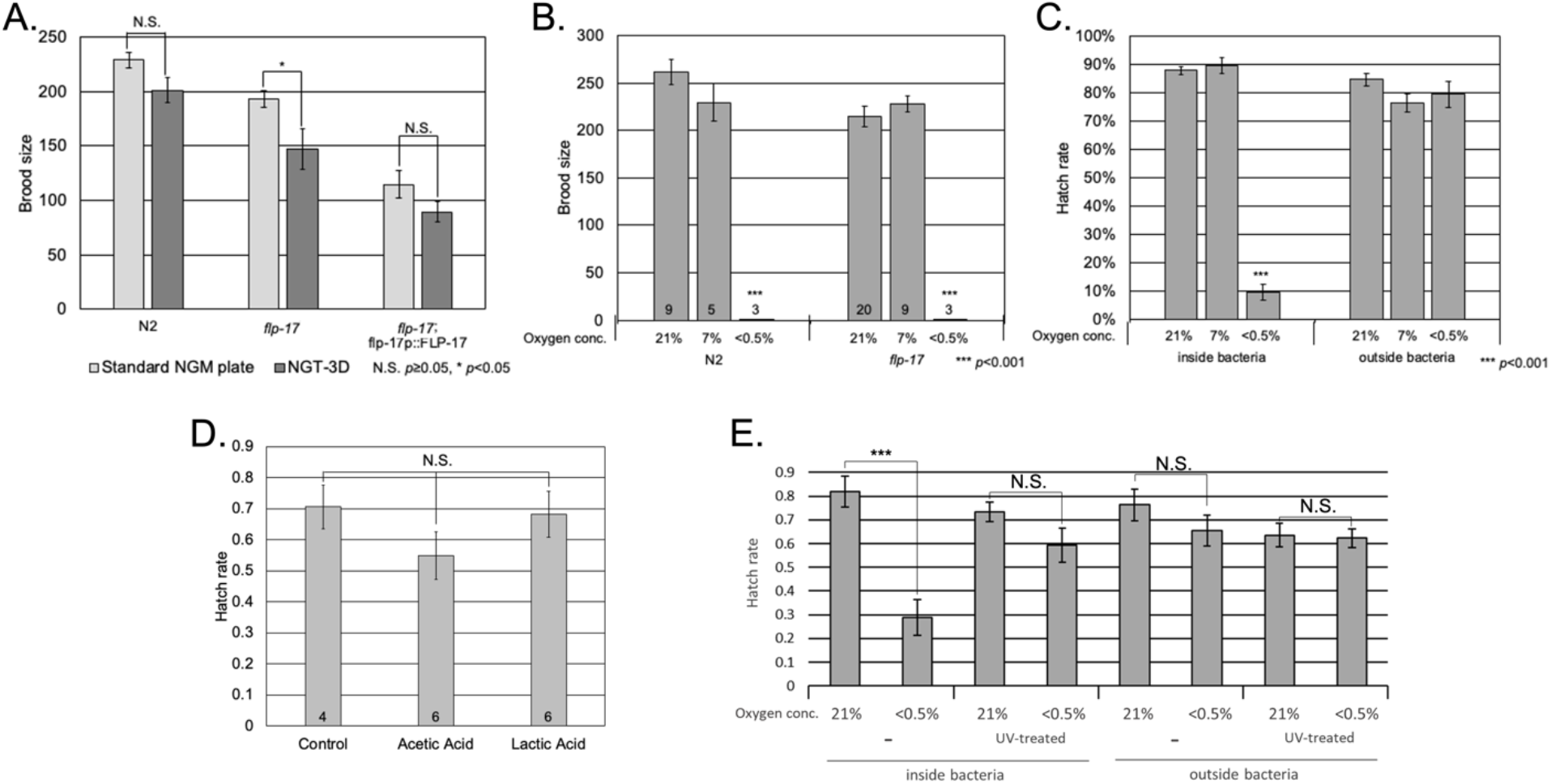
Oviposition behavior increases reproductive fitness by displacing embryos away from toxic low oxygen environments. (A) Differences in brood size in *flp-17* mutants in 2D NGM and NGT-3D. Number of NGM plate or NGT-3D indicated in the bars. (B) Brood size in low oxygen conditions. Number of trials indicated in the bars. (C) Egg viability in low oxygen inside and outside of the bacterial lawn. Number of trials indicated in the bars. (D) Influence of acetic acid or lactic acid on embryo viability. (E) Influence of UV-killed bacteria lawns on embryo viability in hypoxic conditions. Error bars indicate standard error. Statistical significance by student’s T-test indicated. * indicates p < 0.05; ** indicates p < 0.01; *** indicates p < 0.001; NS indicates not significant.

**Figure 7.**
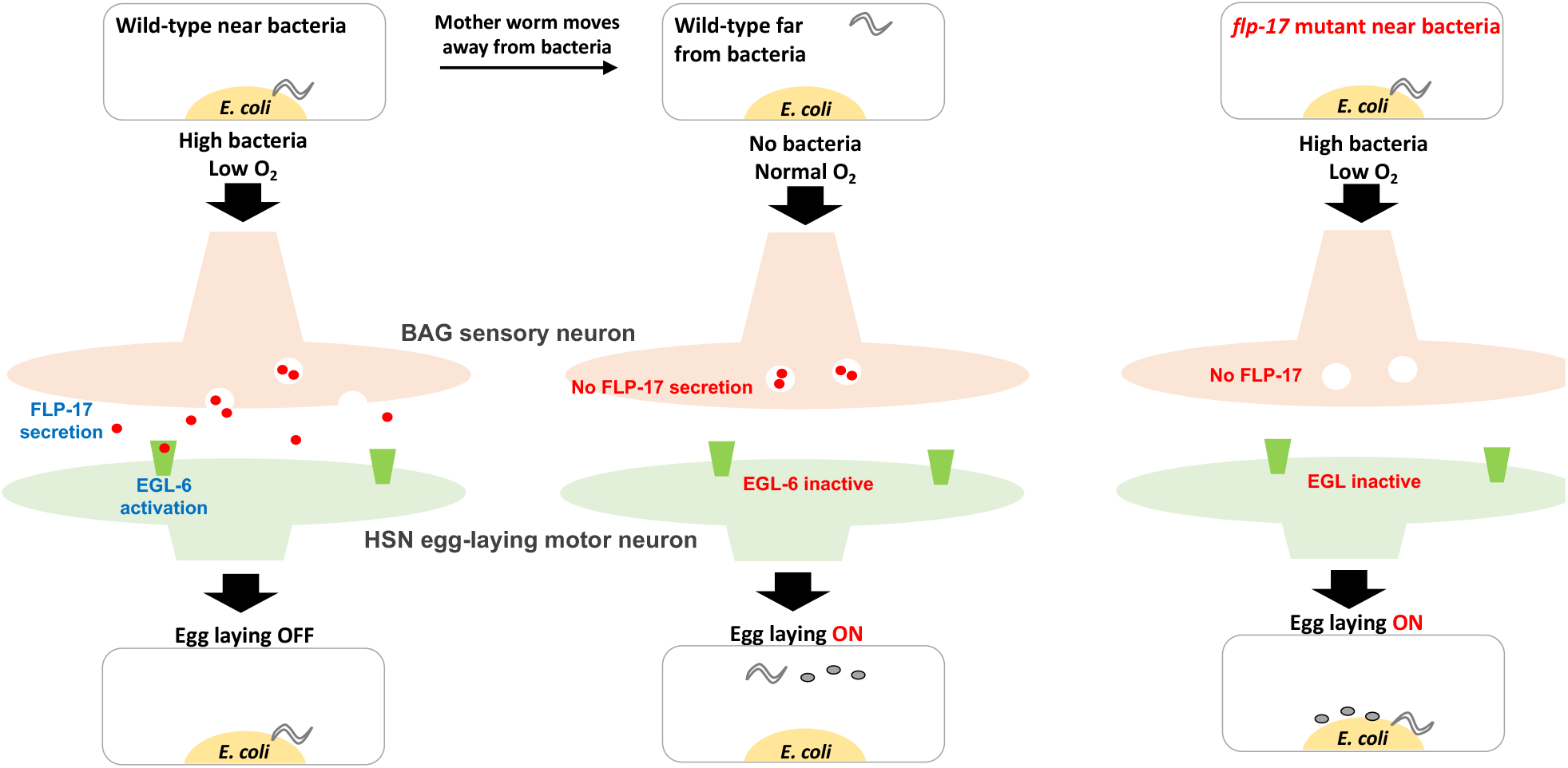
Model of oviposition behavior circuitry.

### Oviposition behavior increases reproductive fitness

Why would such an oxygen-dependent egg laying circuitry exist, and what conditions would allow the adaptation of such a behavior? Maternal behaviors have a direct effect on the reproductive fitness of the mother’s offspring, and convergent evolution of parental behaviors, including oviposition behavior, are observed throughout the animal kingdom^39,40^. We investigated whether *C. elegans* oviposition behavior confers reproductive benefits to mothers by comparing the brood sizes of wild-type N2 and *flp-17* mothers cultivated in 2D and 3D. Although the brood size of N2 mothers is not significantly different in 2D compared to 3D, we found that *flp-17* mothers that cannot display proper oviposition behavior had a significantly smaller brood size in 3D than in 2D (Fig 5A). This defect was restored in the *flp-17* rescue strain (Fig 5A). Although the transgenic rescue strain had a smaller brood size overall regardless of the culture condition, the difference between 2D and 3D was abolished, confirming that oviposition behavior is required for normal brood sizes in 3D. Thus, oviposition behavior confers reproductive fitness benefits to mothers in the 3D environment.

Since *flp-17* mothers that lay eggs near the bacterial colony have lower fitness, we presumed that conditions near the bacteria were toxic for the embryo or young larvae. Since hypoxia is a cue for oviposition behavior (Fig 3), we tested whether low oxygen conditions in 2D resulted in smaller brood size. Surprisingly, there was no significant change in brood size at 7% O2 concentration in both N2 and *flp-17* mutants (Fig 4B). We reasoned that towards the center of the 3D bacteria colony, the O_2_ concentration could reach much lower levels due to *E. coli* aerobic respiration. When we lowered the O_2_ concentration to <0.5%, worms ceased laying eggs, as reported previously^41^. Instead, we placed bleached eggs inside and outside the bacterial lawn and observed embryo viability at different O_2_ levels. At 21% and 7% O_2_, most eggs hatched and there was no difference between eggs inside and outside the lawn. However, at <0.5%, more than 90% of the eggs inside the lawn did not hatch, whereas no effect was observed in hatching outside the lawn.

Embryonic lethality inside the bacteria in low oxygen conditions can be a direct result of hypoxic conditions inside the bacterial lawn or an indirect result of fermentation by the facultative anaerobe *E. coli*. Indeed, fermentative *E. coli* produce copious amounts of lactic acid and acetic acid in anaerobic conditions among other metabolites^42^. Although such high levels of lactic acid do not affect *C. elegans* adult survival^43^, we tested embryo viability in high lactic acid and acetic acid. Interestingly, neither metabolite affected the survival of embryos (Fig 6D).

Finally, to test whether hypoxic conditions inside the bacteria caused embryonic death, we placed eggs inside UV-killed bacteria lawns. We found that in low oxygen conditions, embryo survival increased when they were placed in dead bacteria compared to live bacteria lawns (Fig 6E). This implicates hypoxic conditions brought about by bacterial respiration as a major factor in decreasing embryonic viability. Taken together our work demonstrates that *C. elegans* mother oviposition behavior is likely an adaptation to allow the survival of young in hypoxic environments.

## DISCUSSION

Overall, we have elucidated an oxygen-sensing circuitry that controls egg laying behavior to allow an oviposition behavior that increases the survival of the mothers’ offspring in toxic low oxygen environments (Fig 6). We used a 3D culture condition and found that worms can display oviposition behavior where eggs are laid at a distance from where the bacteria is, even though the mother spends most of its time near the bacteria. We showed that this behavior is mediated by oxygen. We demonstrated that FLP-17, solely expressed in the BAG neuron, is required for this behavior, as well as the receptor EGL-6, expressed in the HSN motor neuron that innervates the vulval muscles. Genetic ablation of BAG and HSN neurons also display the same defective oviposition behavior as *flp-17* and *egl-6* mutants, demonstrating that FLP-17 from BAG neurons is acting on EGL-6 in HSN neurons. Finally, we show that defective oviposition behavior results in lower brood size and thus reduces reproductive fitness, and that this is likely due to extremely low oxygen inside the bacterial colony that prevents hatching.

Based on these results, a plausible model is: in normal oxygen conditions, the BAG sensory neuron does not secrete the FLP-17 neuropeptide^27^, HSN neurons remain active, allowing egg laying to occur normally. However, in low oxygen conditions such as in NGB-3D, the BAG sensory neurons respond to low oxygen and secrete FLP-17, activating the EGL-6 receptor in HSN neurons which inhibits the activity of the neuron to prevent egg laying (Fig 6, left; ^28^). As the mother worm moves away from the bacteria, the oxygen levels increase, inhibiting FLP-17 release allowing the mother to lay her eggs (Fig 6, middle). Finally, in the *flp-17* mutants, the absence of FLP-17 allows the HSN neuron to remain active, and the mother lays eggs near the bacteria, exposing her young to toxic hypoxic conditions.

Since parental behaviors have a direct influence on reproductive fitness, the ability to study maternal care in a simple genetic model animal such as *C. elegans* can elucidate the impact that genes and circuitry regulating a behavior have on the ecology and evolution of an animal. Although the nematode does not display a direct care on her young, *C. elegans* mothers show forms of maternal care such as yolk venting and matriphagy that increase the survival of young^16,44^. During matriphagy, mother worms cease laying eggs in response to a bacterial toxin, allowing their young to hatch inside the mothers’ bodies which provide nutrition for the young to consume and overcome the toxicity^16^. Similarly, we find that the mother worms sense a dangerous combination of hypoxia and bacterial growth and respond by pausing egg laying until she reaches a safer level of oxygen to lay her young. This oxygen-dependent oviposition behavior increases the mother’s reproductive fitness in low oxygen 3D environments similar to the environments that *C. elegans* may be exposed to in nature.

Four neurons in *C. elegans* are known to sense oxygen levels: URX, AQR, PQR, and BAG^45^. The first three are known to sense increases in oxygen levels, whereas BAG neurons are known to sense decreases^32^. The type of soluble guanylyl cyclases expressed in these neurons are thought to account for this difference: URX, AQR, and PQR express *gcy-35* and *gcy-36*, whereas BAG expresses *gcy-33*^*32*^. The BAG sensory neuron is known to coordinate intestinal fat accumulation by sensing low oxygen via GCY-33^27^. This activates the BAG neuron, resulting in FLP-17 release and subsequent deactivation of the URX neuron by the EGL-6 receptor^27^. In this study we have shown that FLP-17 from BAG is required for proper oviposition behavior in 3D and in a low O_2_ environment. It would be interesting to see whether low oxygen is sensed by GCY-33 or another oxygen sensing molecule in the BAG neuron to promote oviposition behavior in NGB-3D or low oxygen environments. Moreover, oxygen’s role in activating the HSN motor neuron and vulva muscle contraction, and whether it would be EGL-6-dependent is unclear. Further studies would be needed to elucidate these questions. The BAG neuron is also known for detecting CO_2_, and increased CO_2_ has been shown to increase the incidence of food leaving behavior^46^. However, we found that higher CO_2_ levels did not produce the oviposition behavior observed in our study.

It is intriguing that the FMRFamide-like neuropeptide FLP-17 precursor peptide shows similarity with gonadotropin-inhibitory hormone (GnIH) precursor peptide that also has a major role in inhibiting reproductive activity in birds and mammals^47,48^. GnIH release from hypothalamic neurons in the brain can block gonadotropin release in the anterior pituitary to affect reproduction in vertebrates^47^. Thus, a role for GnIH peptides in reproductive behaviors is conserved in *C. elegans*.

Although the current study focuses on the location of eggs, oviposition behavior in 3D environments is actually a combination of behaviors. First, the adult worms spread the bacteria around the colony in a sprawling fashion, an observation also made by a previous study^49^. Second, the mother hermaphrodites move away from the bacteria where they normally prefer to stay. Third, they lay eggs and then eventually return to the bacteria. What prompts the worms to move away from the bacteria is not clear. *C. elegans* displays a bacteria-dependent foraging behavior called roaming behavior that is regulated by the neuropeptide PDF-1 and its cognate receptor PDFR-1^50^. To see if this ligand-receptor is involved, we examined oviposition behavior and 3D reproductive fitness of mutants of *pdf-1* and *pdfr-1*. Although *pdf-1* mutants showed normal oviposition behavior and reproductive fitness, *pdfr-1* mutants were defective for both (Fig S3), indicating that roaming may be required for proper oviposition behavior.

In this study we have identified an oxygen-dependent oviposition behavior in the N2 Bristol strain of *C. elegans*. However, *C. elegans* can be found throughout the world, and many different strains have been isolated^38^. Variation in behaviors such as exploratory behavior^51^, oxygen sensing^37,52^, and matricidal egg laying^53^ have been described across the *C. elegans* species. Similarly, we found that genetically diverse natural strains of *C. elegans* display a large degree of variation in oviposition behavior with two of these strains displaying significant differences. These results support the idea that evolution may be shaping a maternal behavior through genetic variation similar to that observed in the timing of egg laying in the European great tit^4^. What ecological factors may be regulating these natural differences? Although the 11 wild strains we tested have been isolated from geographically diverse areas, it is difficult to derive any clear conclusions about the ecology of these strains since most were found in vegetative or fruit compost and soil including the two behaviorally different strains^14,54^. What genetic factors may be regulating these natural differences? We compared the *flp-17* and *egl-6* genes in N2 and the JU258 and JT11398 strains but we did not find any sequence changes. Thus, another gene or genes may account for the differences in oviposition behavior.

Behavior studies in genetic models such as *C. elegans*, the fruit fly and the mouse, have elucidated the role of genes, neurons and circuitry in animal behavior. However, the impact of these behaviors and genes on the survival and reproduction of the animal in its true ecological environment can be difficult to ascertain. Our NGB-3D environment certainly cannot replicate *C. elegans*’ actual habitats in nature, but it adds an important level of complexity that has not been tested before. The fact that we were able to reveal a novel behavior that can affect reproductive fitness demonstrates the further need for such studies. In addition, our study suggests that the plasticity of behaviors such as egg laying is a necessity for success in diverse and potentially dangerous habitats, implying that such behaviors are an adaptive product of evolution.

## ACKNOWLEDGMENTS

We thank the Caenorhabditis Genetic Center, CeNDR and Niels Ringstad for strains and Supriya Srinivasan for resources. We thank Wormbase, CeNDR, CeNGEN for information and other resources. We thank Rocel A. Indong for critical reading of the manuscript. This research was funded by the National Research Foundation of Korea grant number 2021R1A2C101178312 (J.I.L.).

## AUTHOR CONTRIBUTIONS

Conceptualization: T.Y.L and J.I.L; Methodology: T.Y.L and J.I.L; Software: K-h.Y.; Investigation: T.Y.L and E.C.; Formal Analysis: T.Y.L, E.C., and J.I.L; Writing – Original Draft: J.I.L, T.Y.L., and K-h.Y., Project Administration: J.I.L.; Funding Acquisition: J.I.L.

## STAR METHODS

### Key resources table

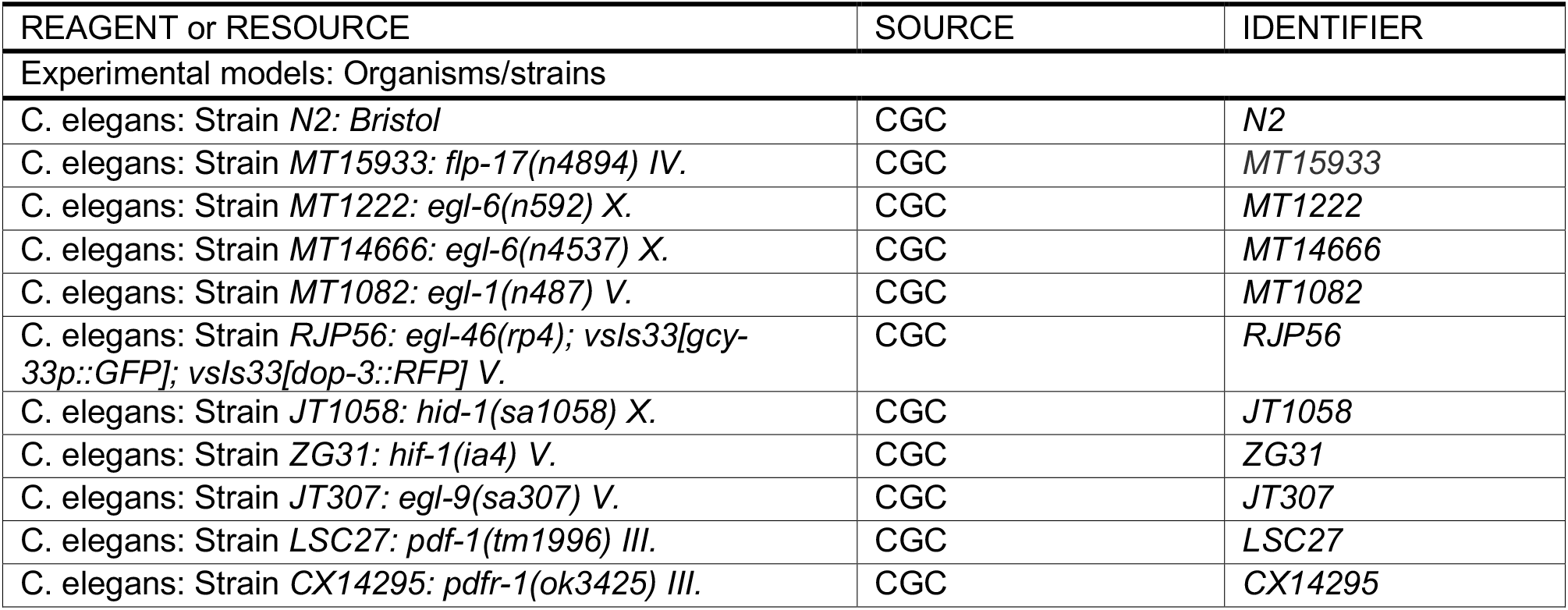
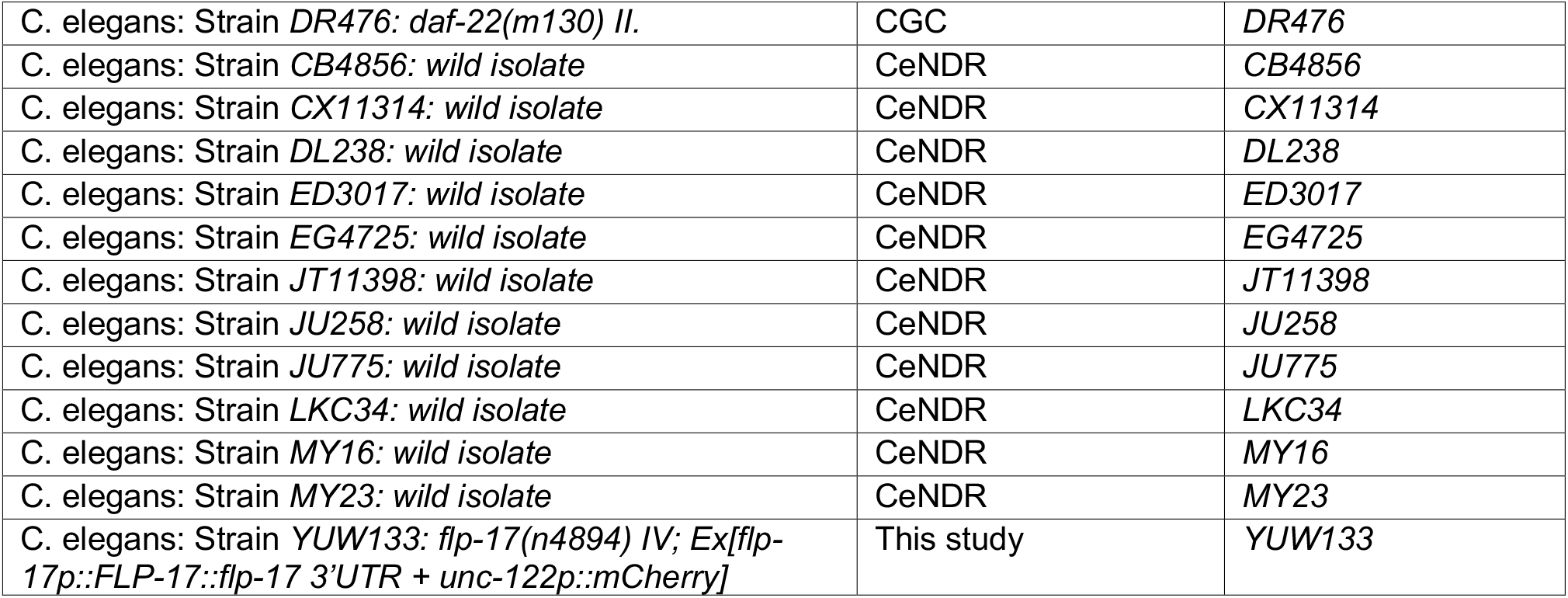

### Resource availability

Further information and requests for resources and reagents should be directed to and will be fulfilled by the lead contact Jin I. Lee

### Materials availability

All unique reagents generated in this study are available from the lead contact.

### Experimental model and subject details

All the *C. elegans* strains are maintained on the standard NGM plate with seeded *E. coli* OP50 at 20°C using standard methods^55^. A detailed list of all strains used is provided in the key resources table.

### Method details

#### Imaging

All photos and movies were captured using Nikon DS-Fi2 camera mounted on Nikon SMZ18 stereomicroscope, with NIS-Elements Basic Research software.

#### Nematode Growth Media plates

Standard techniques were used to make NGM agar plates^55^ except that Difco Granulated Agar was used as the solidifying agent (20 g in 1 liter media). OP50 dot plates were prepared according to a previous study^17,18^. Briefly, a platinum wire was dipped in an overnight *E. coli* OP50 culture in LB media (Difco), and was used to place 5 evenly spaced dots of *E. coli* across a 5.5 cm fresh NGM plate. Plates were incubated overnight at 37°C resulting in dense colonies with diameters between 1-2 mm.

#### Nematode Growth Bottle-3D

NGB-3D was prepared according to a previous study^18^. Briefly, autoclaved NGM media was cooled to 40°C, and 30ml diluted bacteria was added directly to the media. Diluted bacteria was prepared by serial dilution of overnight culture of *E. coli* OP50 to 1×10^−8^ with 0.85% NaCl solution. After mixing, 65ml of the media was poured into a 25 cm^2^ cell culture flask (SPL) and left for at least 1 week at room temperature for bacterial growth.

#### Nematode Growth Tube-3D

NGT-3D was prepared according to a previous study^17,18^ and was prepared and seeded the same way as NGB-3D. 6.5ml of media was poured into 10 ml clear plastic test tubes (SciLab) and was briefly left to harden. A layer of NGM agar without OP50 was poured on top to prevent colonies forming too close to the surface and left for at least 1 week at room temperature for bacterial growth. To check general oxygen levels in NGT-3D, thioglycolate media was used instead of NGM media (29.5 g BBL Fluid Thioglycolate medium, 4.25g Difco Granulated Agar for 1L media).

#### Recording 3D Behavior

Well-fed L4 stage worms were picked, and bacteria on the body was cleaned off by placing the worm on a non-seeded plate for several minutes and repeated twice. The cleaned mother worm was placed in one of the culture chambers with a platinum wire pick. For 3D behavior observation, ten bacteria-free worms were initially placed on the agar surface of NGB-3D, incubated for 24hr and behavior was observed under the microscope.

#### Quantifying 3D behavior

To measure bacterial colony radius and spread bacteria (aka. sprawl) radius, the ImageJ Polygon selection tool was used. The border line of the colony and sprawl was selected using the polygon selection tool, and Feret diameter was measured. Most of the bacterial colonies have a consistent shape, but the sprawl can be quite varied in shape. Thus, separate formulas were used for collecting colony and sprawl radius. For determining the radius of the bacterial colony, the following geometric mean formula was used:

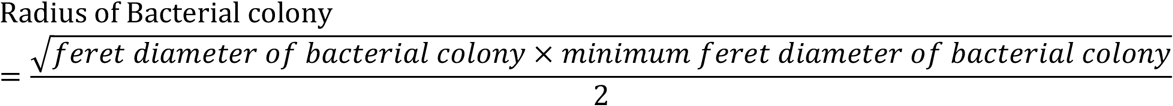

To determine the sprawl radius, the following minimum ferret diameter formula was used:

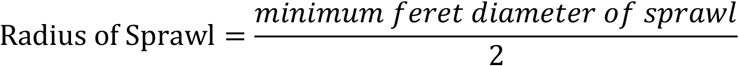

To determine the distance between the tip of the mother worm nose and the center of the bacterial colony (ie. worm distance), a continuous 200 second movie after 24hr incubation in NGB-3D was recorded. Worm distance then was collected every 10 seconds using the ImageJ Straight tool. To determine the distance between egg location and the center of the bacterial colony (aka. egg distance), first, the assumption was made that the bacterial colony is a perfect sphere was made. Based on the assumption, three layers of images in the Z-axis were captured based on the focus of the colony: a top, middle and bottom layer. The clearest egg image was then found and the distance between the egg and the center of the colony was measured using the ImageJ straight tool. The egg distance data was then calculated using the Pythagorean theorem:

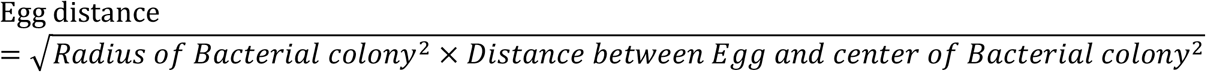

To calculate the oviposition ratio, the following formula was used with the worm distance and egg distance data:

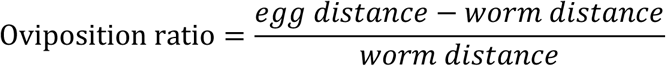

This ratio shows how far the worms lay eggs from where they are normally located. Since more than one mother worm can occupy the same colony, it was not possible to determine exactly which mother had laid each egg. Thus, oviposition ratio was calculated with an average egg distance for all eggs at each colony for each individual mother worm at that colony.

For navigation behavior, a single bacteria-free worm was placed on an OP50 dot plate or on the agar surface of NGB-3D, and observed under the microscope. A 10-minute video was recorded immediately after the worm first contacted the bacteria, and navigation behavior was analyzed based on the 10 minute movie as previously performed^19^ and adjusted for this study. Briefly, worm locomotion was categorized into 7 groups. All the behaviors were categorized at every 1 second interval by the researcher. A straight movement longer than an adult *C. elegans* body size was scored as long forward or long reverse movement, and movement shorter than their body size was scored as short forward or short reverse movement. Omega turns were visually identified by the head nearly touching the tail or a reorientation of >90° within a single head swing. A short movement in the opposite direction less than 1 second or a complete stop for more than 1 second was scored as a pause. Any moment in which the worm either moved out of frame or behind the colony or otherwise was not able to be observed, was scored as other.

#### 2D egg laying observations

To test the influence of gas levels, five bacteria-free worms were placed on a dot plate and incubated for 24hr. To test at different gas levels, plates were placed in a modular incubator chamber (STEMCELL, Cat. # 27310) with an adjustable intake valve for gas delivery. Industrial-use gas sensor indicator was placed inside to monitor gas levels (oxygen indicator: SENKO, SP2nd and RS Korea, Oxygen Gas Detector, S20082109; carbon dioxide indicator: FORENSICS DETECTORS, FD-90A). Plates were imaged after the incubation period. To determine lawn radius, the geometric mean formula was used based on ferret diameter measured by the ImageJ polygon selection tool. Egg distance was measured by the ImageJ straight tool.

#### Molecular biology and obtaining transgenic animals

Promoters and genes were generated using standard molecular biology and PCR techniques. *C. elegans* genomic DNA was obtained from N2 Bristol worm lysates and purified using a tissue DNA purification kit (Labopass). Construction of the *flp-17p::FLP-17* plasmid was based on a similar reagent from a previous study^27^, where 3165 bp fragment from *C. elegans* genomic DNA containing 1347 bp of the *flp-17* upstream promoter region, and the *flp-17* gene was used. This fragment was obtained by PCR using Q5 High-Fidelity DNA Polymerase (New England Biolabs), the forward primer [5’-GAA ACC CGG CCG ATG AAG G-3’] and the reverse primer [5’-CAA ATA GGC AGC GTT TCA TTG-3’] with *C. elegans* genomic DNA as the template. The fragment was subcloned using TOPcloner blunt Kit (Enzynomics) into a pTOP blunt V2 vector, and purified using PureLink miniprep kit (Invitrogen). Transgenic *C. elegans* were obtained by microinjection into the *C. elegans* germline followed by visual selection of transgenic animals under fluorescence. For microinjections, 70 ng/μl of the target plasmid was co-injected with 30ng/μl of *unc-122p::mCherry* as a co-injection marker and 20 ng/μl of an empty pUC19 vector to maintain a total injection mix concentration of 120 ng/μl. In each case, several stable transgenic lines were generated. A single line (YUW133) was selected for experimentation based on consistency of expression and transmission rate.

#### Brood size assay

A single cleaned L4 worm was placed on either an NGM plate or NGT-3D with a platinum wire pick, and incubated for 96 h. Brood size was counted as previously described^17^. Briefly, NGT-3D and the NGM plate were melted in an 88°C water bath and poured into 9 cm plates, where dead but intact worms could be scored. Only adult, L4, and L3 worms were counted to ensure that only the F1 generation (and the P0 worm) is counted to avoid confusion with the younger F2 generation. Student’s t-test was used to determine significance.

#### Egg viability assay

Eggs were collected by treating gravid adult worms with bleach solution as described previously^55^. After washing, 100 eggs were placed on the plate seeded with 20 µl of an overnight culture of OP50 *E. coli* either directly in or outside the bacterial lawn and incubated for 48hr in the modular incubator chamber. Hatch rate was calculated by the following formula:

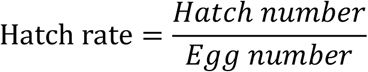

For acetic acid and lactic acid experiments, acetic acid or lactic acid (10 mM final concentration) were added to NGM media minus cholesterol. 100 eggs were placed in the center of the 9 cm plate and an overnight *E. coli* OP50 culture in LB media was carefully spread in a thin line shape surrounding the eggs with a 2 cm gap and incubated for 48hr.

For the UV exposure experiment, the *E. coli* OP50 plates were exposed to 6mJ/cm UV light for 20 min with Spectrolinker™ XL-1000 Series UV Crosslinker. The plates were exposed with lids open, along with the inner side of the lid being exposed. After 20 min exposure, the plates were placed in 20°C incubator to cool down before using. To confirm whether UV exposure completely killed *E. coli* OP50, the 20 µl *E. coli* OP50 lawn was entirely scratched off and streaked using a 10ul loop onto an antibiotic free LB agar plate, incubated 37°C overnight along with a non-UV killed bacteria streak plate as a control group. The UV exposed *E. coli* OP50 plates showed no growth confirming that UV exposure completely killed the bacteria.

## SUPPLEMENTAL INFORMATION TITLES AND LEGENDS

**Figure S1.**
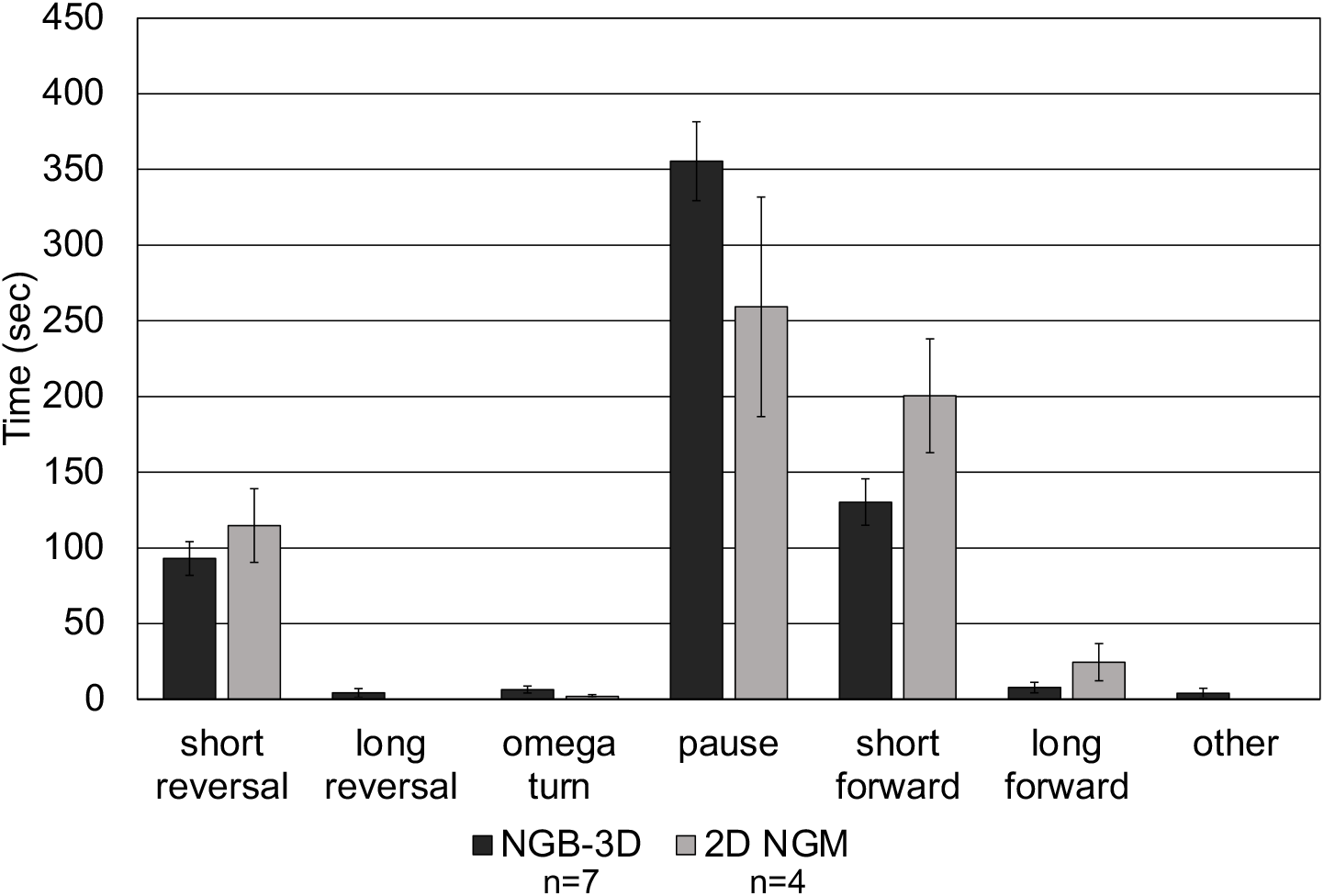
Comparison of navigation movement behavior in 2D and 3D. Mother worms were placed on either NGB-3D or 2D NGM patch plates with bacteria lawn diameters of 3-5 mm. Immediately upon contacting the bacteria, navigation movement was recorded for 10 min. Number of trials is indicated above the bars.

**Figure S2.**
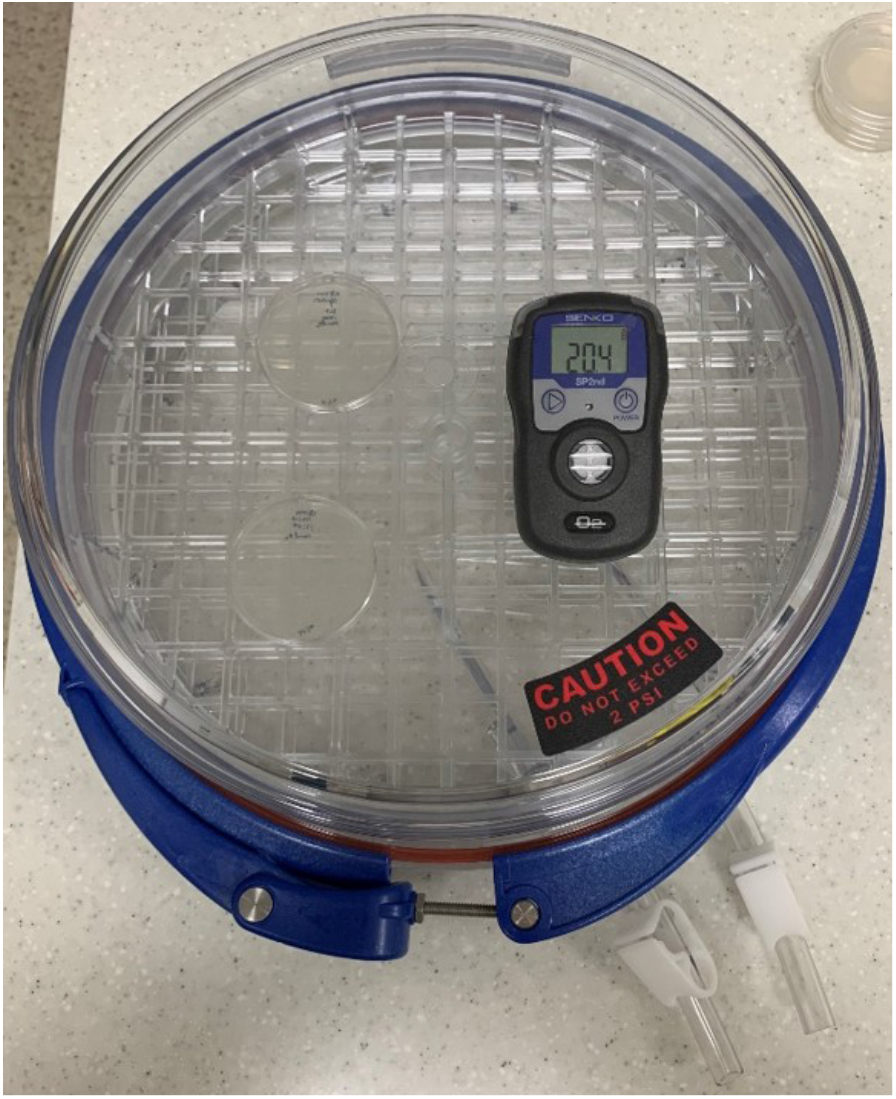
Modular incubator chamber with industrial gas sensor for assessing egg laying in different gas concentrations.

**Figure S3.**
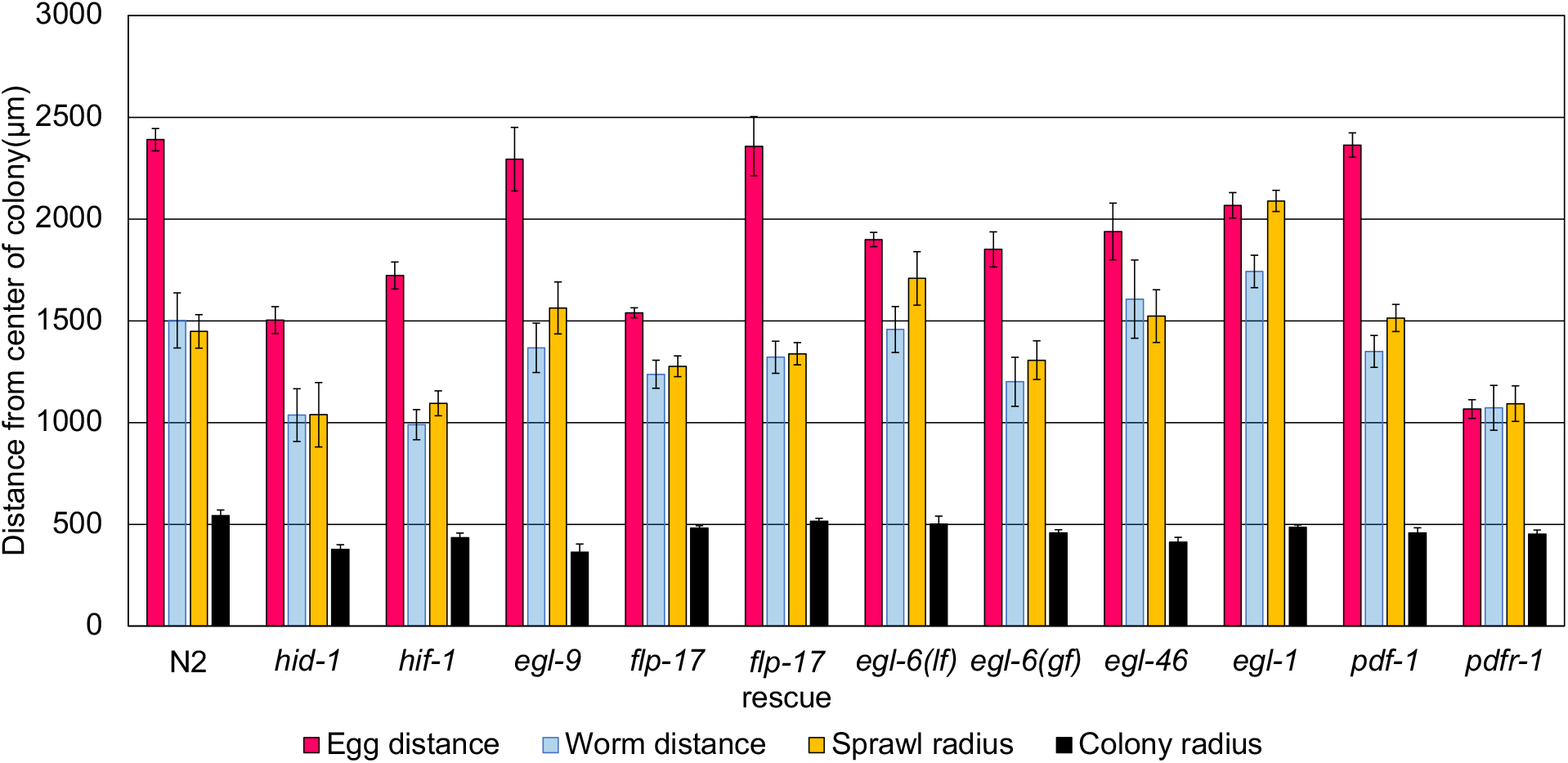
3D oviposition behavior in different mutant strains. Error bars indicate standard error.

**Figure S4.**
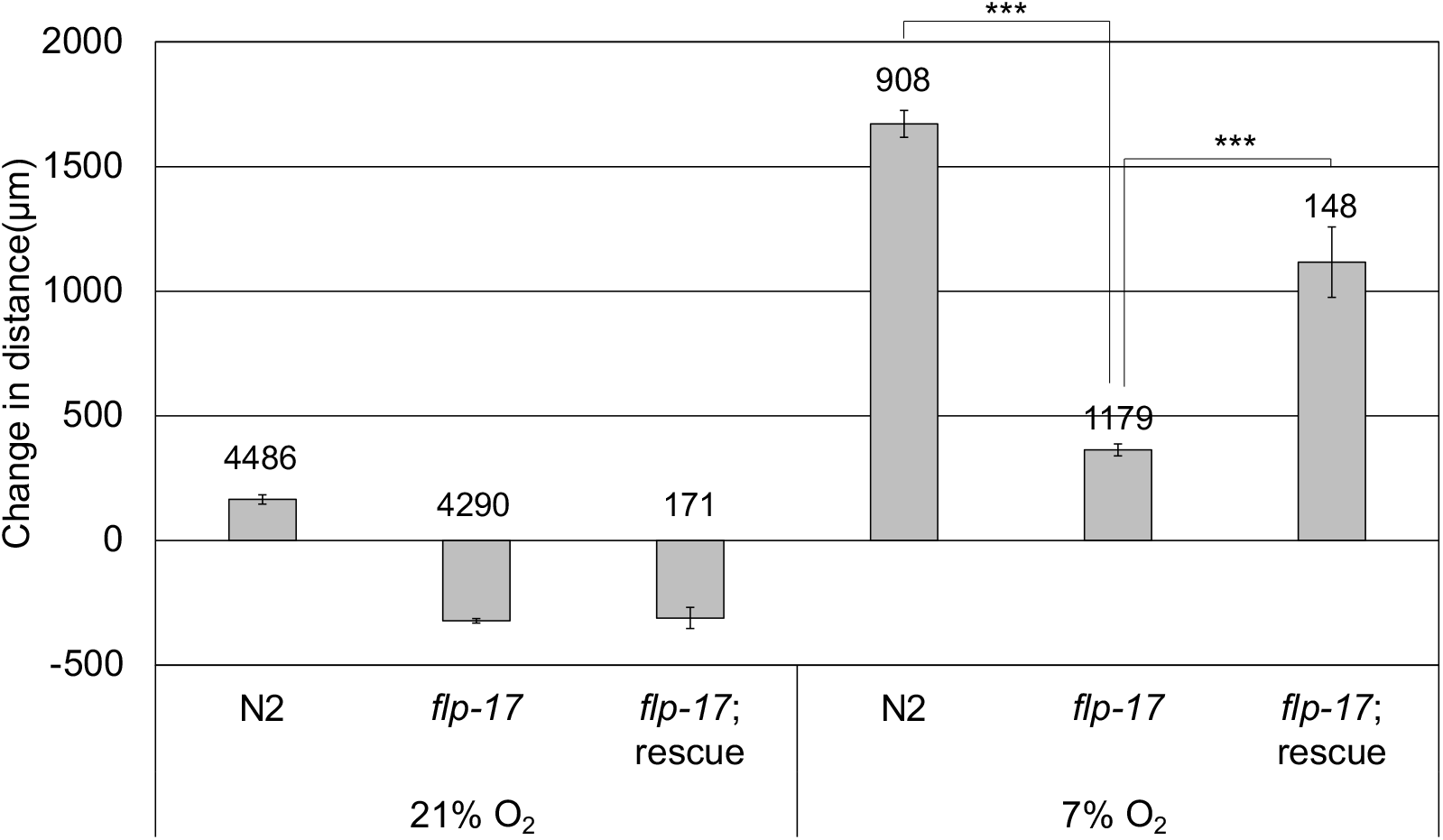
Distance from lawn edge of total laid eggs in 2D at normal and low oxygen levels. Total laid eggs is all eggs laid in all experiments regardless of bacterial lawn. Number of eggs counted is indicated above the bars. Error bars indicate standard error. Significance is determined by Student’s T-test. * indicates p < 0.05. *** indicates p < 0.001.

**Fig. S5.**
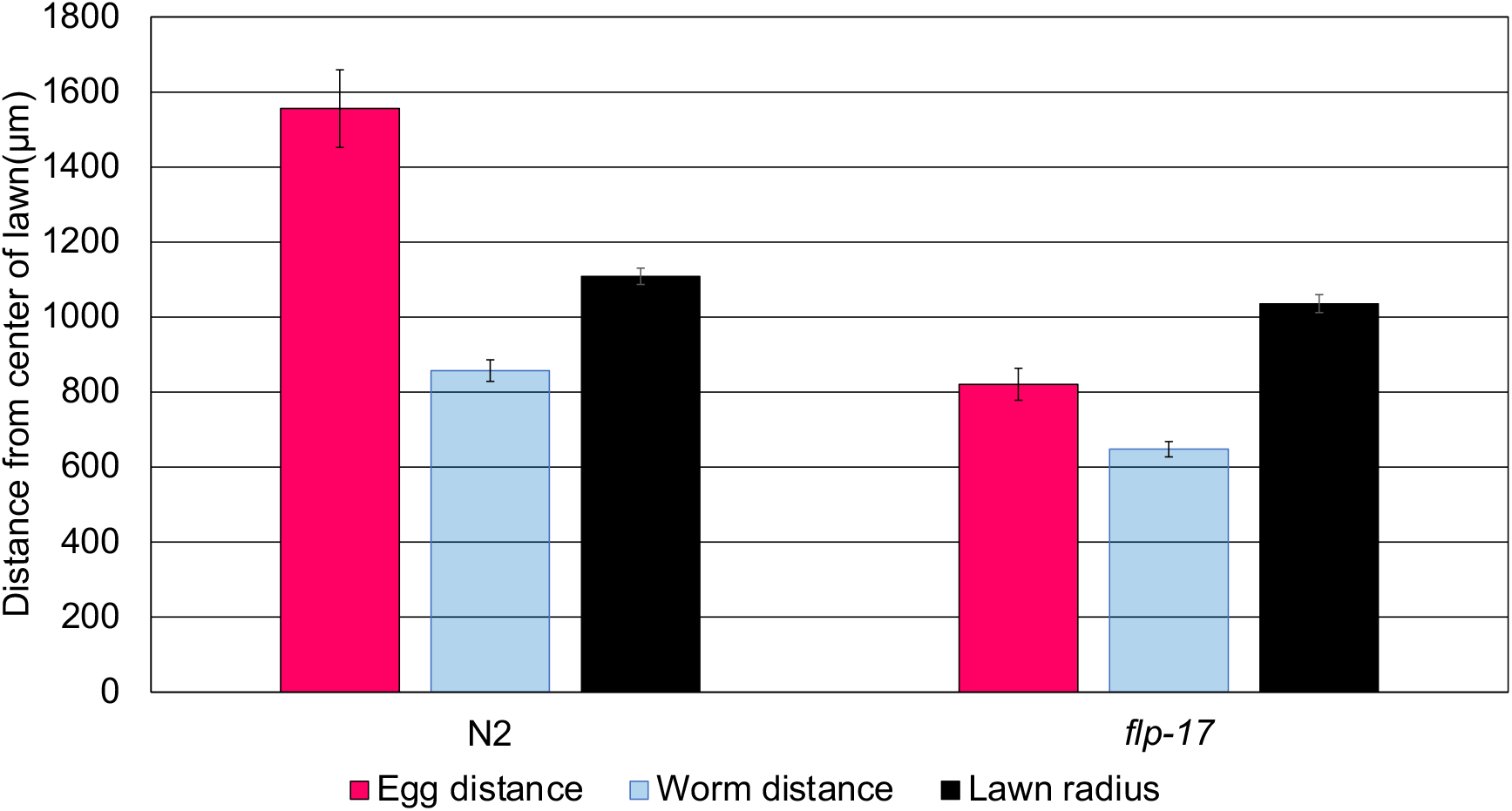
Egg distance, worm distance and lawn radius in 2D NGM dot plate at 21% oxygen. Error bars indicate standard error.

Video S1. 24 hr time lapse video of an N2 adult hermaphrodite *C. elegans* in a bacterial colony in NGB-3D. Note the spreading of the bacterial sprawl, and the eggs laid away from the bacteria. A second adult worm joins after 20 hours. Bar indicates 1 mm.

Video S2. N2 adult hermaphrodite mother *C. elegans* laying eggs in a 2D NGM plate in normal speed. Red circle indicates moment of egg laying. Bar indicates 1 mm.

Video S3. N2 adult hermaphrodite mother *C. elegans* laying eggs in NGB-3D sped up 4x. Red circle indicates moment of egg laying. Bar indicates 1 mm.

Video S4. *flp-17* mutant adult hermaphrodite mother *C. elegans* laying eggs in a 2D NGM plate sped up 5x. Arrow indicates head of mother worm. Red circle indicates moment of egg laying. Bar indicates 1 mm.

## REFERENCES

1. Royle, N.J., Russell, A.F., and Wilson, A.J. (2014). The evolution of flexible parenting. Science 345, 776–781. 10.1126/science.1253294.

2. Westneat, D.F., Hatch, M.I., Wetzel, D.P., and Ensminger, A.L. (2011). Individual variation in parental care reaction norms: integration of personality and plasticity. Am Nat 178, 652–667. 10.1086/662173.

3. Ghalambor, C.K., Peluc, S.I., and Martin, T.E. (2013). Plasticity of parental care under the risk of predation: how much should parents reduce care? Biol Lett 9, 20130154. 10.1098/rsbl.2013.0154.

4. Nussey, D.H., Postma, E., Gienapp, P., and Visser, M.E. (2005). Selection on heritable phenotypic plasticity in a wild bird population. Science 310, 304–306. 10.1126/science.1117004.

5. Fang, Y.Y., Yamaguchi, T., Song, S.C., Tritsch, N.X., and Lin, D. (2018). A Hypothalamic Midbrain Pathway Essential for Driving Maternal Behaviors. Neuron 98, 192–207 e110. 10.1016/j.neuron.2018.02.019.

6. Wu, Z., Autry, A.E., Bergan, J.F., Watabe-Uchida, M., and Dulac, C.G. (2014). Galanin neurons in the medial preoptic area govern parental behaviour. Nature 509, 325–330. 10.1038/nature13307.

7. Kohl, J., Babayan, B.M., Rubinstein, N.D., Autry, A.E., Marin-Rodriguez, B., Kapoor, V., Miyamishi, K., Zweifel, L.S., Luo, L., Uchida, N., and Dulac, C. (2018). Functional circuit architecture underlying parental behaviour. Nature 556, 326–331. 10.1038/s41586-018-0027-0.

8. Yang, C.H., Belawat, P., Hafen, E., Jan, L.Y., and Jan, Y.N. (2008). Drosophila egg-laying site selection as a system to study simple decision-making processes. Science 319, 1679–1683. 10.1126/science.1151842.

9. Takamura, T., and Fuyama, Y. (1980). Behavior genetics of choice of oviposition sites in Drosophila melanogaster. I. Genetic variability and analysis of behavior. Behav Genet 10, 105–120. 10.1007/BF01067322.

10. Dweck, H.K., Ebrahim, S.A., Kromann, S., Bown, D., Hillbur, Y., Sachse, S., Hansson, B.S., and Stensmyr, M.C. (2013). Olfactory preference for egg laying on citrus substrates in Drosophila. Curr Biol 23, 2472–2480. 10.1016/j.cub.2013.10.047.

11. Mansourian, S., Enjin, A., Jirle, E.V., Ramesh, V., Rehermann, G., Becher, P.G., Pool, J.E., and Stensmyr, M.C. (2018). Wild African Drosophila melanogaster Are Seasonal Specialists on Marula Fruit. Curr Biol 28, 3960–3968 e3963. 10.1016/j.cub.2018.10.033.

12. Iliff, A.J., and Xu, X.Z.S. (2020). C. elegans: a sensible model for sensory biology. J Neurogenet 34, 347–350. 10.1080/01677063.2020.1823386.

13. Schafer, W.F. (2006). Genetics of egg-laying in worms. Annu Rev Genet 40, 487–509. 10.1146/annurev.genet.40.110405.090527.

14. Barriere, A., and Felix, M.A. (2005). High local genetic diversity and low outcrossing rate in Caenorhabditis elegans natural populations. Curr Biol 15, 1176–1184. 10.1016/j.cub.2005.06.022.

15. Quach, K.T., and Chalasani, S.H. (2022). Flexible reprogramming of Pristionchus pacificus motivation for attacking Caenorhabditis elegans in predator-prey competition. Curr Biol 32, 1675–1688 e1677. 10.1016/j.cub.2022.02.033.

16. Yoon, K.H., Lee, T.Y., Moon, J.H., Choi, S.Y., Choi, Y.J., Mitchell, R.J., and Il Lee, J. (2020). Consumption of Oleic Acid During Matriphagy in Free-Living Nematodes Alleviates the Toxic Effects of the Bacterial Metabolite Violacein. Sci Rep 10, 8087. 10.1038/s41598-020-64953-x.

17. Lee, T.Y., Yoon, K.H., and Lee, J.I. (2016). NGT-3D: a simple nematode cultivation system to study Caenorhabditis elegans biology in 3D. Biol Open 5, 529–534. 10.1242/bio.015743.

18. Lee, T.Y., Yoon, K.H., and Lee, J.I. (2016). Cultivation of Caenorhabditis elegans in Three Dimensions in the Laboratory. J Vis Exp. 10.3791/55048.

19. Gray, J.M., Hill, J.J., and Bargmann, C.I. (2005). A circuit for navigation in Caenorhabditis elegans. Proc Natl Acad Sci U S A 102, 3184–3191. 10.1073/pnas.0409009101.

20. Brewer, J.H. (1940). Clear liquid mediums for the aerobic cultivation of anaerobes. JAMA 115, 598–600. 10.1001/jama.1940.72810340001009.

21. Twigg, R. (1945). Oxidation-Reduction Aspects of Resazurin. Nature 155, 401-402. doi.org/10.1038/155401a0.

22. Sharabi, K., Charar, C., Friedman, N., Mizrahi, I., Zaslaver, A., Sznajder, J.I., and Gruenbaum, Y. (2014). The response to high CO2 levels requires the neuropeptide secretion component HID-1 to promote pumping inhibition. PLoS Genet 10, e1004529. 10.1371/journal.pgen.1004529.

23. Jiang, H., Guo, R., and Powell-Coffman, J.A. (2001). The Caenorhabditis elegans hif-1 gene encodes a bHLH-PAS protein that is required for adaptation to hypoxia. Proc Natl Acad Sci U S A 98, 7916–7921. 10.1073/pnas.141234698.

24. Epstein, A.C., Gleadle, J.M., McNeill, L.A., Hewitson, K.S., O’Rourke, J., Mole, D.R., Mukherji, M., Metzen, E., Wilson, M.I., Dhanda, A., et al. (2001). C. elegans EGL-9 and mammalian homologs define a family of dioxygenases that regulate HIF by prolyl hydroxylation. Cell 107, 43–54. 10.1016/s0092-8674(01)00507-4.

25. Chang, A.J., and Bargmann, C.I. (2008). Hypoxia and the HIF-1 transcriptional pathway reorganize a neuronal circuit for oxygen-dependent behavior in Caenorhabditis elegans. Proc Natl Acad Sci U S A 105, 7321–7326. 10.1073/pnas.0802164105.

26. McVeigh, P., Geary, T.G., Marks, N.J., and Maule, A.G. (2006). The FLP-side of nematodes. Trends Parasitol 22, 385–396. 10.1016/j.pt.2006.06.010.

27. Hussey, R., Littlejohn, N.K., Witham, E., Vanstrum, E., Mesgarzadeh, J., Ratanpal, H., and Srinivasan, S. (2018). Oxygen-sensing neurons reciprocally regulate peripheral lipid metabolism via neuropeptide signaling in Caenorhabditis elegans. PLoS Genet 14, e1007305. 10.1371/journal.pgen.1007305.

28. Ringstad, N., Abe, N., and Horvitz, H.R. (2009). Ligand-gated chloride channels are receptors for biogenic amines in C. elegans. Science 325, 96–100. 10.1126/science.1169243.

29. Kim, K., and Li, C. (2004). Expression and regulation of an FMRFamide-related neuropeptide gene family in Caenorhabditis elegans. J Comp Neurol 475, 540–550. 10.1002/cne.20189.

30. Bretscher, A.J., Busch, K.E., and de Bono, M. (2008). A carbon dioxide avoidance behavior is integrated with responses to ambient oxygen and food in Caenorhabditis elegans. Proc Natl Acad Sci U S A 105, 8044–8049. 10.1073/pnas.0707607105.

31. Hallem, E.A., and Sternberg, P.W. (2008). Acute carbon dioxide avoidance in Caenorhabditis elegans. Proc Natl Acad Sci U S A 105, 8038–8043. 10.1073/pnas.0707469105.

32. Zimmer, M., Gray, J.M., Pokala, N., Chang, A.J., Karow, D.S., Marletta, M.A., Hudson, M.L., Morton, D.B., Chronis, N., and Bargmann, C.I. (2009). Neurons detect increases and decreases in oxygen levels using distinct guanylate cyclases. Neuron 61, 865–879. 10.1016/j.neuron.2009.02.013.

33. Rojo Romanos, T., Petersen, J.G., Riveiro, A.R., and Pocock, R. (2015). A novel role for the zinc-finger transcription factor EGL-46 in the differentiation of gas-sensing neurons in Caenorhabditis elegans. Genetics 199, 157–163. 10.1534/genetics.114.172049.

34. Godini, R., Langebeck-Jensen, K., and Pocock, R. (2020). A single amino acid change in the EGL-46 transcription factor causes defects in BAG neuron specification. MicroPubl Biol 2020. 10.17912/micropub.biology.000224.

35. Conradt, B., and Horvitz, H.R. (1998). The C. elegans protein EGL-1 is required for programmed cell death and interacts with the Bcl-2-like protein CED-9. Cell 93, 519–529. 10.1016/s0092-8674(00)81182-4.

36. Nehme, R., and Conradt, B. (2008). egl-1: a key activator of apoptotic cell death in C. elegans. Oncogene 27 Suppl 1, S30–40. 10.1038/onc.2009.41.

37. McGrath, P.T., Rockman, M.V., Zimmer, M., Jang, H., Macosko, E.Z., Kruglyak, L., and Bargmann, C.I. (2009). Quantitative mapping of a digenic behavioral trait implicates globin variation in C. elegans sensory behaviors. Neuron 61, 692–699. 10.1016/j.neuron.2009.02.012.

38. Cook, D.E., Zdraljevic, S., Roberts, J.P., and Andersen, E.C. (2017). CeNDR, the Caenorhabditis elegans natural diversity resource. Nucleic Acids Res 45, D650–D657. 10.1093/nar/gkw893.

39. Fischer, E.K., Roland, A.B., Moskowitz, N.A., Vidoudez, C., Ranaivorazo, N., Tapia, E.E., Trauger, S.A., Vences, M., Coloma, L.A., and O’Connell, L.A. (2019). Mechanisms of Convergent Egg Provisioning in Poison Frogs. Curr Biol 29, 4145–4151 e4143. 10.1016/j.cub.2019.10.032.

40. Fox, C.W., Wagner, J.D., Cline, S., Thomas, F.A., and Messina, F.J. (2009). Genetic architecture underlying convergent evolution of egg-laying behavior in a seed-feeding beetle. Genetica 136, 179–187. 10.1007/s10709-008-9334-y.

41. Nystul, T.G., and Roth, M.B. (2004). Carbon monoxide-induced suspended animation protects against hypoxic damage in Caenorhabditis elegans. Proc Natl Acad Sci U S A 101, 9133–9136. 10.1073/pnas.0403312101.

42. Clark, D.P. (1989). The fermentation pathways of Escherichia coli. FEMS Microbiol Rev 5, 223–234. 10.1016/0168-6445(89)90033-8.

43. Gomez, F., Monsalve, G.C., Tse, V., Saiki, R., Weng, E., Lee, L., Srinivasan, C., Frand, A.R., and Clarke, C.F. (2012). Delayed accumulation of intestinal coliform bacteria enhances life span and stress resistance in Caenorhabditis elegans fed respiratory deficient E. coli. BMC Microbiol 12, 300. 10.1186/1471-2180-12-300.

44. Kern, C.C., Townsend, S., Salzmann, A., Rendell, N.B., Taylor, G.W., Comisel, R.M., Foukas, L.C., Bahler, J., and Gems, D. (2021). C. elegans feed yolk to their young in a form of primitive lactation. Nat Commun 12, 5801. 10.1038/s41467-021-25821-y.

45. Carrillo, M.A., and Hallem, E.A. (2015). Gas sensing in nematodes. Mol Neurobiol 51, 919–931. 10.1007/s12035-014-8748-z.

46. Milward, K., Busch, K.E., Murphy, R.J., de Bono, M., and Olofsson, B. (2011). Neuronal and molecular substrates for optimal foraging in Caenorhabditis elegans. Proc Natl Acad Sci U S A 108, 20672–20677. 10.1073/pnas.1106134109.

47. Tsutsui, K., and Ubuka, T. (2021). Gonadotropin-inhibitory hormone (GnIH): A new key neurohormone controlling reproductive physiology and behavior. Front Neuroendocrinol 61, 100900. 10.1016/j.yfrne.2021.100900.

48. Ubuka, T., and Tsutsui, K. (2018). Comparative and Evolutionary Aspects of Gonadotropin-Inhibitory Hormone and FMRFamide-Like Peptide Systems. Front Neurosci 12, 747. 10.3389/fnins.2018.00747.

49. Thutupalli, S., Uppaluri, S., Constable, G.W., Levin, S.A., Stone, H.A., Tarnita, C.E., and Brangwynne, C.P. (2017). Farming and public goods production in Caenorhabditis elegans populations. Proc Natl Acad Sci U S A 114, 2289–2294. 10.1073/pnas.1608961114.

50. Flavell, S.W., Pokala, N., Macosko, E.Z., Albrecht, D.R., Larsch, J., and Bargmann, C.I. (2013). Serotonin and the neuropeptide PDF initiate and extend opposing behavioral states in C. elegans. Cell 154, 1023–1035. 10.1016/j.cell.2013.08.001.

51. Bendesky, A., Tsunozaki, M., Rockman, M.V., Kruglyak, L., and Bargmann, C.I. (2011). Catecholamine receptor polymorphisms affect decision-making in C. elegans. Nature 472, 313–318. 10.1038/nature09821.

52. Persson, A., Gross, E., Laurent, P., Busch, K.E., Bretes, H., and de Bono, M. (2009). Natural variation in a neural globin tunes oxygen sensing in wild Caenorhabditis elegans. Nature 458, 1030–1033. 10.1038/nature07820.

53. Vigne, P., Gimond, C., Ferrari, C., Vielle, A., Hallin, J., Pino-Querido, A., El Mouridi, S., Mignerot, L., Frokjaer-Jensen, C., Boulin, T., et al. (2021). A single-nucleotide change underlies the genetic assimilation of a plastic trait. Sci Adv 7. 10.1126/sciadv.abd9941.

54. Andersen, E.C., Gerke, J.P., Shapiro, J.A., Crissman, J.R., Ghosh, R., Bloom, J.S., Felix, M.A., and Kruglyak, L. (2012). Chromosome-scale selective sweeps shape Caenorhabditis elegans genomic diversity. Nat Genet 44, 285–290. 10.1038/ng.1050.

55. Lewis, J.A., and Fleming, J.T. (1995). Basic culture methods. Methods Cell Biol 48, 3–29.

