## Supplementary figures and images for "The neuropeptide FLP-17 regulates an oviposition behavior in the nematode *Caenorhabditis elegans* that increases maternal reproductive fitness in low oxygen environments"

### Fig S1

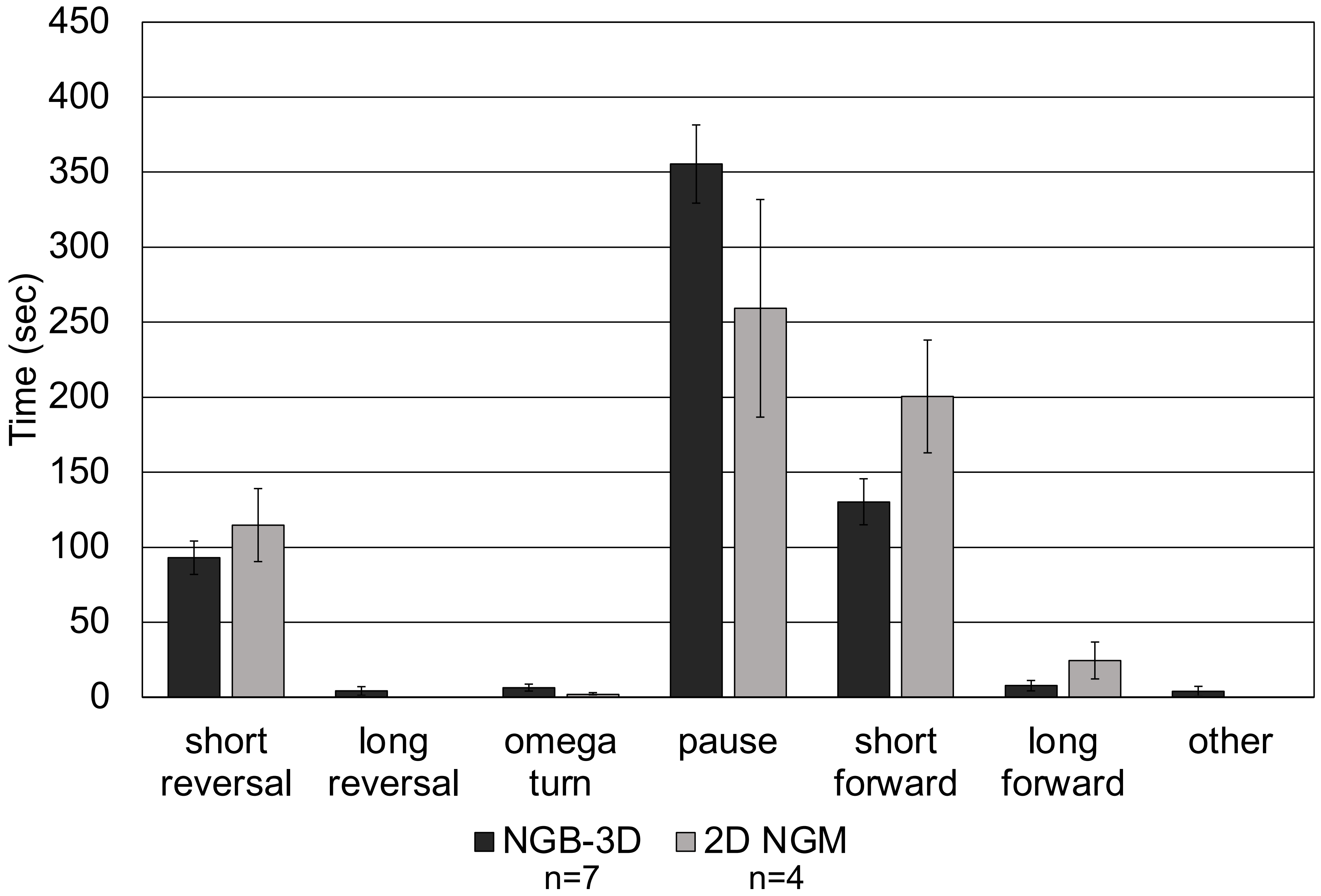

### Fig S2

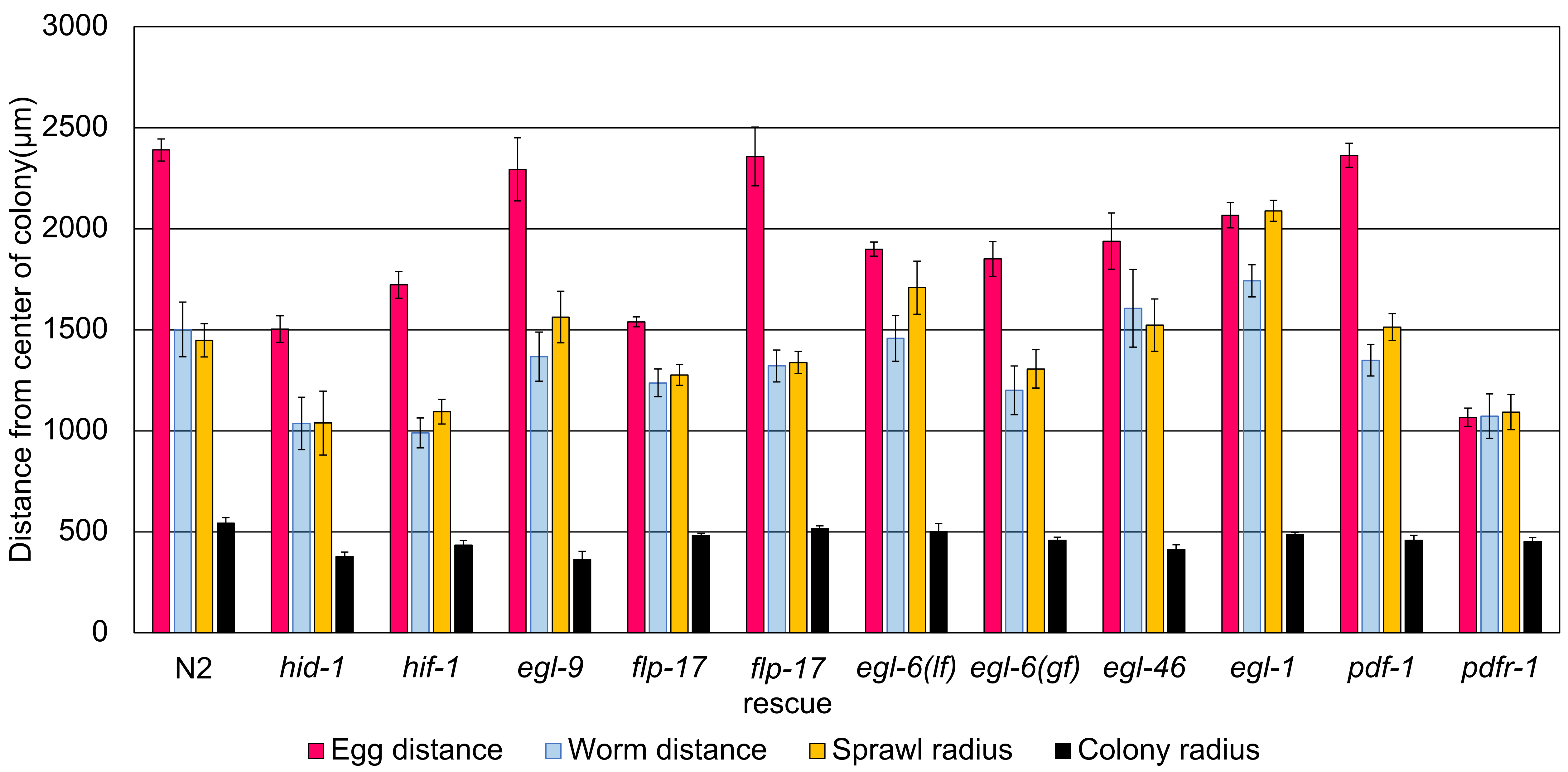

### Fig S3

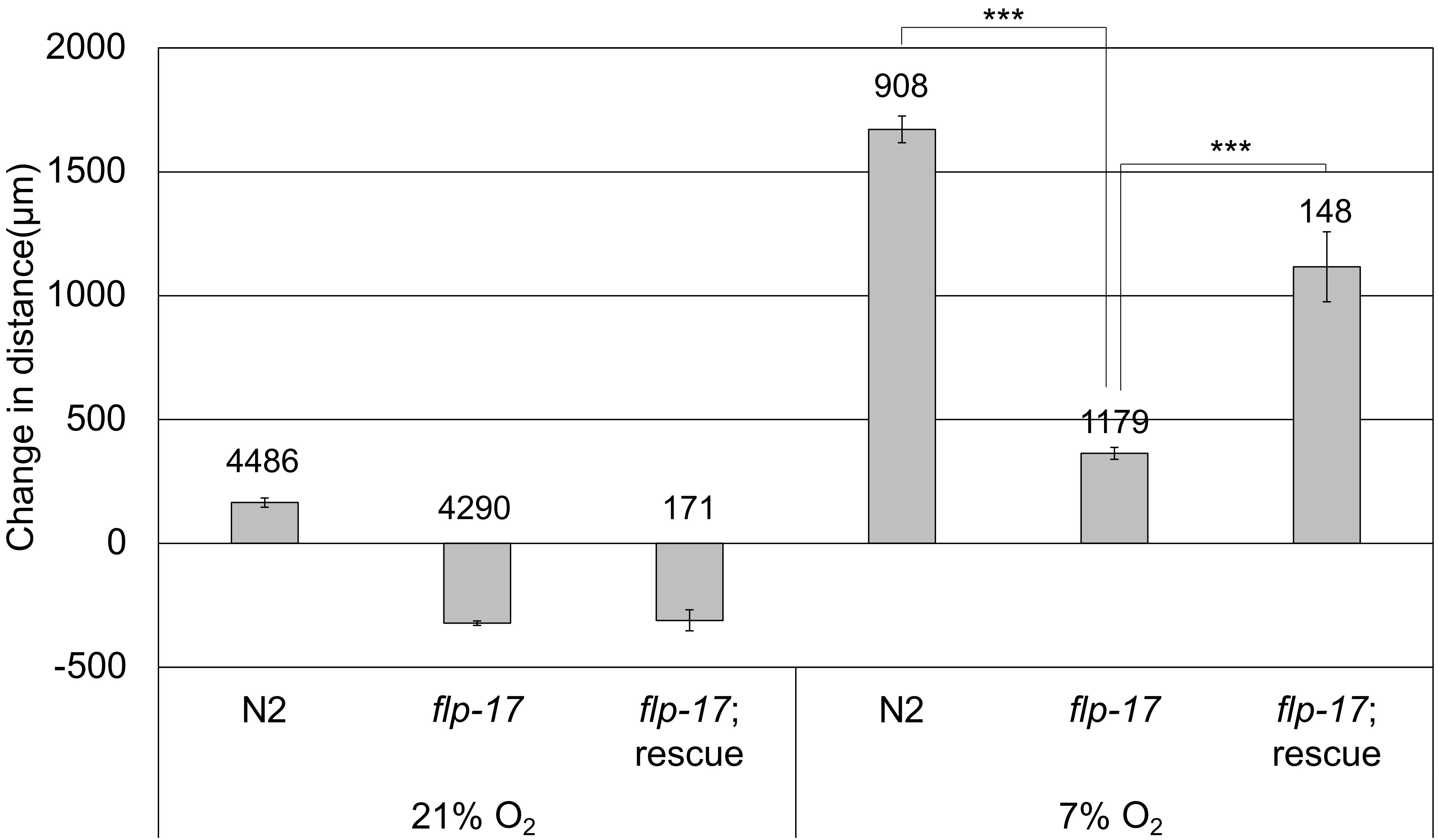

### Fig S4

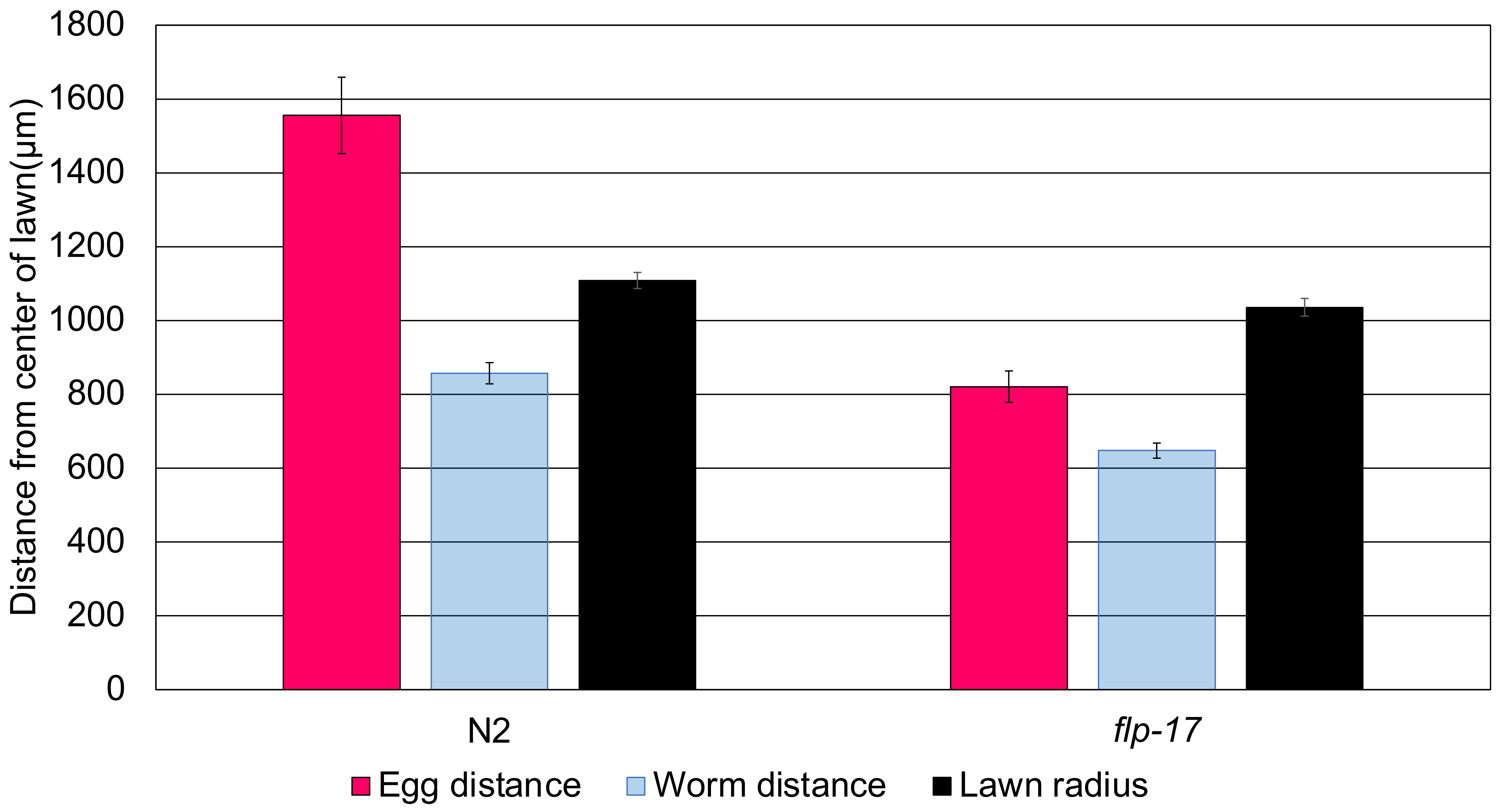
